# Phosphorylation of disordered proteins tunes local and global intramolecular interactions

**DOI:** 10.1101/2024.06.10.598315

**Authors:** Emery T. Usher, Martin J. Fossat, Alex S. Holehouse

**Affiliations:** Department of Biochemistry and Molecular Biophysics, Washington University School of Medicine, St. Louis, MO, USA; Center for Biomolecular Condensates (CBC), Washington University in St. Louis, St. Louis, MO, USA; Department of Biological Physics, Max Planck Institute of Immunobiology and Epigenetics, Freiburg im Breisgau, Germany

## Abstract

Protein post-translational modifications, such as phosphorylation, are important regulatory signals for diverse cellular functions. In particular, intrinsically disordered protein regions (IDRs) are subject to phosphorylation as a means to modulate their interactions and functions. Toward understanding the relationship between phosphorylation in IDRs and specific functional outcomes, we must consider how phosphorylation affects the IDR conformational ensemble. Various experimental techniques are suited to interrogate the features of IDR ensembles; molecular simulations can provide complementary insights and even illuminate ensemble features that may be experimentally inaccessible. Therefore, we sought to expand the tools available to study phosphorylated IDRs by all-atom Monte Carlo simulations. To this end, we implemented parameters for phosphoserine (pSer) and phosphothreonine (pThr) into the OPLS version of the continuum solvent model, ABSINTH, and assessed their performance in all-atom simulations compared to published findings. We simulated short (< 20 residues) and long (> 80 residues) phospho-IDRs that, collectively, survey both local and global phosphorylation-induced changes to the ensemble. Our simulations of four well-studied phospho-IDRs show near-quantitative agreement with published findings for these systems via metrics including changes to radius of gyration, transient helicity, and persistence length. We also leveraged the inherent advantage of sequence control in molecular simulations to explore the conformational effects of diverse combinations of phospho-sites in two multi-phosphorylated IDRs. Our results support and expand on prior observations that connect phosphorylation to changes in the IDR conformational ensemble. Herein, we describe phosphorylation as a means to alter sequence chemistry, net charge and charge patterning, and intramolecular interactions, which can collectively modulate the local and global IDR ensemble features.

**SIGNIFICANCE:** Spatially and temporally controlled phosphorylation in disordered protein regions is critical to many facets of protein function and broader cellular health. Intrinsically disordered protein regions (IDRs) are overrepresented as targets of phosphorylation, but the structural and functional consequences of such modifications remain elusive for many systems. Toward rigorous modeling of phosphorylated IDRs using all-atom simulations, we present new parameters for phosphoserine and phosphothreonine for the ABSINTH implicit solvent paradigm. Through the study of four example phospho-IDRs, we demonstrate excellent agreement between our phospho-IDR simulations and published datasets.

## INTRODUCTION

Protein phosphorylation is an integral regulatory signal for many cellular processes including transcription, cell cycle control, and signal transduction (1–7). Intrinsically disordered protein regions (IDRs) are overrepresented as targets of such modifications owing to their relative enrichment in modifiable residues and accessible surface area (8–10). Moreover, temporally and spatially specific modification of IDRs grants reversible control over the conformational ensemble and intermolecular interactions that support protein function (11–18). As such, there is a need for high-resolution, quantitative descriptions of IDR ensembles and how phosphorylation may bias the conformations accessed by the protein.

Solution-state experimental methods, such as nuclear magnetic resonance (NMR) spectroscopy, small-angle X-ray scattering (SAXS), and Förster Resonance Energy Transfer (FRET) can collectively provide ensemble-level and atomistic details of IDR conformations (19–24). Molecular simulations of IDRs are a powerful complement to experiments; simulations enable the atomistic study of the sequence-ensemble relationship of IDRs (25–30). As such, we aim to use simulations to interrogate how changes to IDR sequence chemistry through PTMs alter ensemble behavior.

Molecular dynamics (MD) and Monte Carlo (MC) simulation strategies have seen specific recent developments toward robust and computationally tractable modeling of IDRs. Improvements to force fields, solvent models, and sampling schemes for MD and MC simulations of IDRs enable their quantitative characterization *in silico* (31–33). Many IDR-optimized force fields and common explicit water models are already equipped with parameters for phosphorylated residues and have been used to computationally probe phospho-IDR conformational space (34–40). However, these parameters are not necessarily interoperable across force fields and solvent models. Therefore, we sought to develop and assess parameters for phosphoserine (pSer) and phosphothreonine (pThr) to be used with the OPLS version of the ABSINTH implicit solvent model (*see Methods*).

Here, we report parameters to describe pSer and pThr for ABSINTH-OPLS and demonstrate their application through four case study proteins. Test proteins were selected based on size, number of phosphorylation events, and the reported phosphorylation-induced effects; we ultimately chose two short peptides and two longer IDRs (**Table 1**). A fragment of the sodium-proton exchanger 1 (NHE1, residues 781-796) containing one phospho-site was chosen for its prior characterization by NMR spectroscopy and MD simulations. Hendus-Altenberger and colleagues found that the transient helical character of this peptide is stabilized by N-terminal phosphorylation (41). For our second system, we chose a short peptide from Tau (residues 173 - 183) containing two phospho-sites. Independent studies reported local stiffening of Tau upon phosphorylation based on NMR and FRET experiments and coarse-grained MD simulations (42, 43) making it an ideal candidate for our validation

**Table 1:**
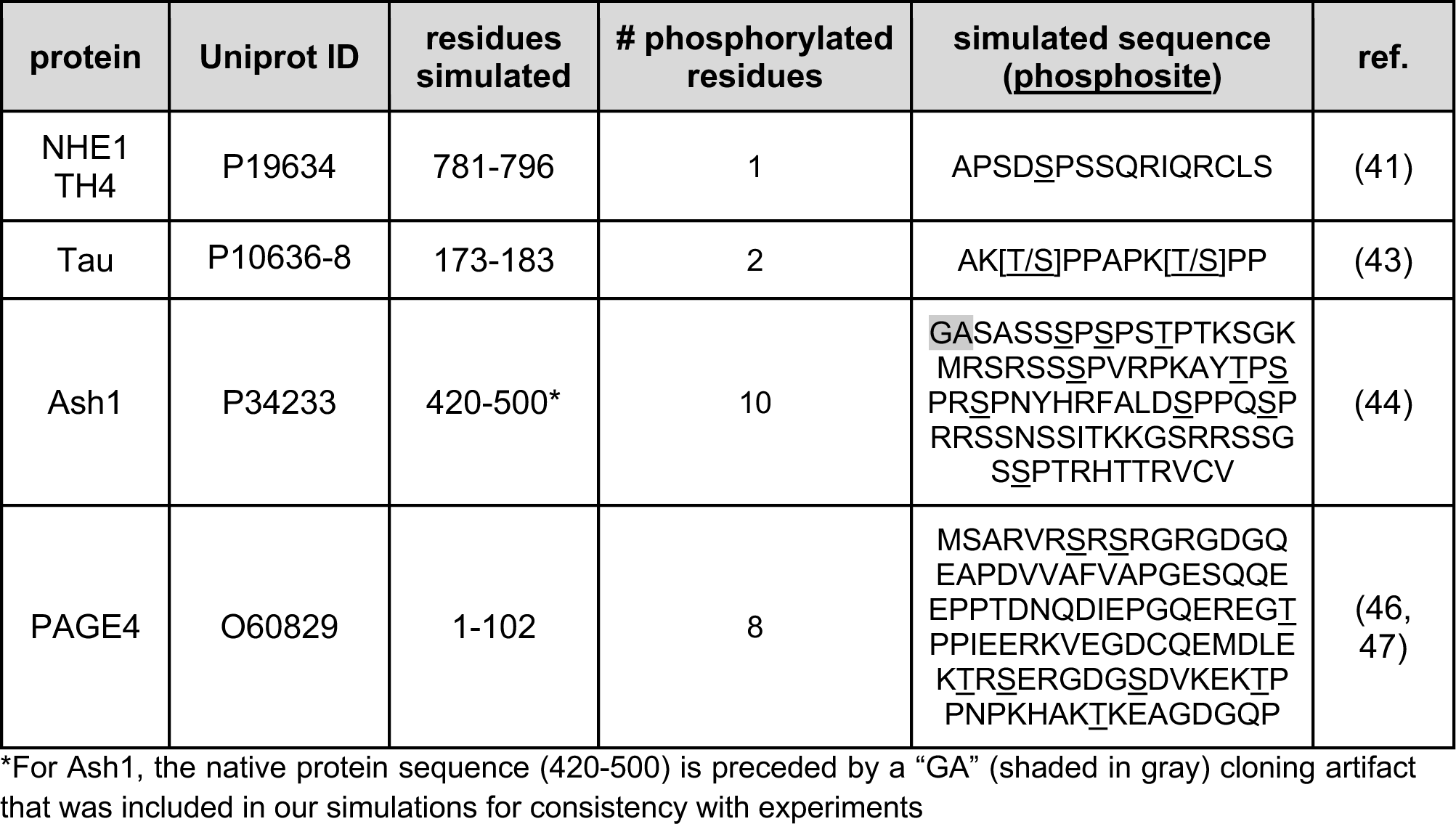
Names, accession numbers, and sequences for phospho-protein simulations.

To assess our pSer/pThr parameters in the context of longer sequences, we selected two well-characterized IDRs that undergo multi-site phosphorylation (**Table 1**). An IDR from the yeast transcription factor Ash1 (residues 420-500) was characterized experimentally in its unmodified and hyper-phosphorylated forms by NMR and SAXS. This study also reported CAMPARI simulations of Ash1 using a phosphomimetic sequence (pSer/pThr represented by glutamate) that were in quantitative agreement with experiments on the true phosphorylated Ash1 IDR (44). We also simulated the Prostate-Associated Gene 4 (PAGE4) protein, which is a transcriptional coactivator upregulated in prostate cancer (45). The full-length PAGE4 has 102 residues, contains no globular domains, and can be phosphorylated up to eight times. Further, the PAGE4 ensemble has been described by experiments and simulations in three distinct phospho-states, which makes it an ideal phospho-IDR for testing our simulation parameters (46–48).

Across the four chosen phospho-IDR systems, our simulations capture the phosphorylation-induced changes to local secondary structure and global ensemble dimensions reported previously by experiments and simulations (41–44, 48) (**Figure 1A**). Acknowledging the reported discrepancies across force fields and solvent models for phosphorylated residues, we do not expect strict quantitative agreement of our ensembles with those from different simulation methods or experimental measurements (34, 35, 49). Rather, we sought to (i) establish a parameter set for phosphorylated residues to work within the CAMPARI-ABSINTH framework; (ii) demonstrate robust sampling of phospho-IDR conformational space; and (iii) understand the applications and limitations of these simulations for well-studied IDR systems using prior experiments and simulations for comparison.

**Figure 1:**
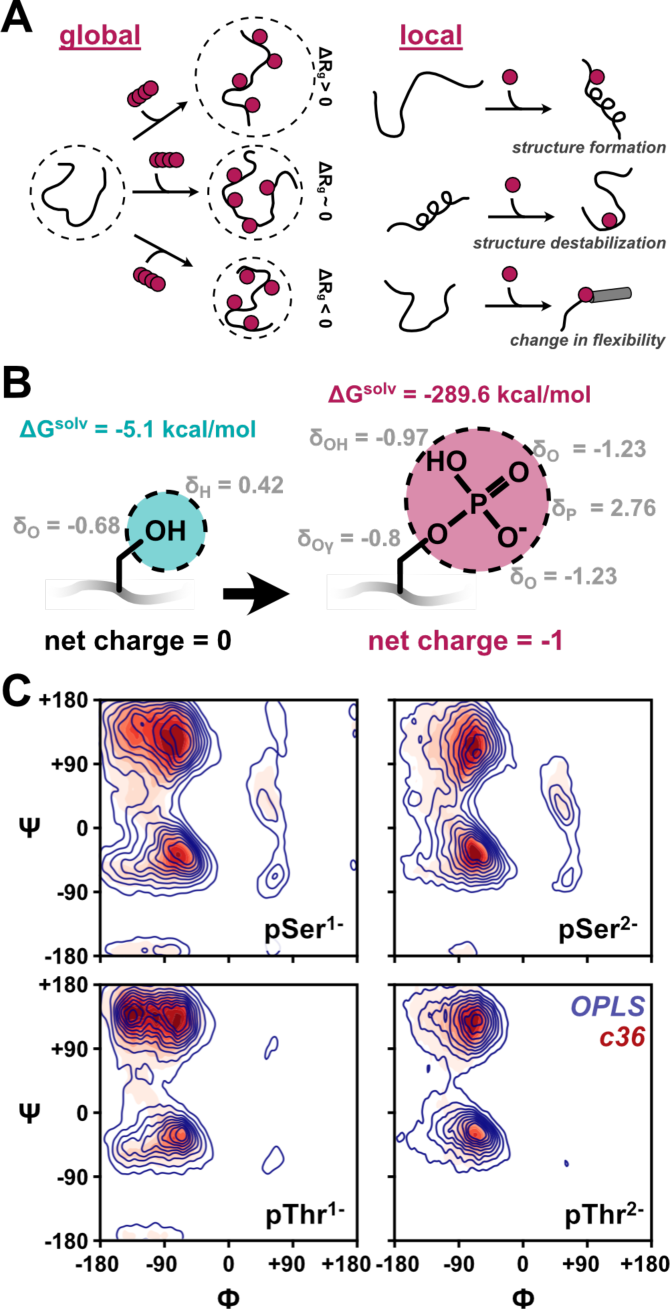
Summary of pSer and pThr parameters and observables used for validation. **(A)** Diagram of possible global and local ensemble effects associated with phosphorylation. **(B)** Free energy of solvation (FOS) values, per-atom partial charges, and side chain connectivity for a hydroxyl group versus a phosphoryl group. **(C)** Ramachandran plots of the phi/psi space sampled by the pSer or pThr residue in 3-mer peptides.

## METHODS

Implicit or continuum solvent models are particularly attractive for simulating IDRs due to the lowered computational expense compared with explicitly modeling each water molecule. To this end, the ABSINTH implicit solvation model was developed with the considerations and challenges of IDR simulations in mind (50). The CAMPARI simulation engine enables Metropolis MC simulations using the ABSINTH implicit solvent paradigm to efficiently explore IDR conformational space (51). Thus, CAMPARI simulations of IDRs using the OPLS-AA force field-derived charge assignments (ABSINTH-OPLS) yield ensemble features that are consistent with those from diverse experimental methods (44, 52–58).

### Phospho-serine and -threonine parameter implementation in CAMPARI

Parameters for phosphoserine (pSer, Sep) and phosphothreonine (pThr, Tpo) were implemented in CAMPARI v3 (https://campari.sourceforge.net/) with parameter sets *abs3.5_opls_phos.prm* or *abs3.5_charmm36_phos.prm* modified to contain the values for free energy of solvation (FOS) and per-atom partial charges for the relevant phosphate charge states (**Table S1**). The FOS for the z = -1 and -2 phosphoryl groups on pSer or pThr were calculated from experimental methods using the protocol described in reference (59). Briefly, we used the Thermodynamic Cycle based on Proton Dissociation (TCPD) method to calculate hydration free energies 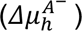 for each pSer or pThr side chain in each relevant charge state based on Equation 1:

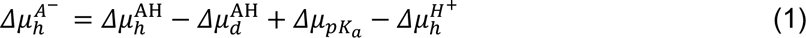

Where 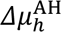 is hydration free energy of the protonated form of the acid, 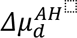 and Δμ*_pka_* are the change in free energy from dissociation of a proton in the gas phase and aqueous phase, respectively. 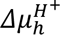 is the free energy of hydration of the proton, which was taken from reference (59). For z = -2, A^-^ refers to the fully deprotonated phosphoryl moiety and AH refers to the singly protonated phosphoryl group. Similarly for z = -1, A^-^ is the singly protonated phosphoryl group and AH is the fully protonated (uncharged) phosphoryl group.

For z = -2 and -1, gas-phase basicities (proton affinities) for phosphate compounds were taken from reference (60). For both pSer and pThr, experimental pKa,2 values from reference (61) were used. For the z = -1 TCPD calculations, we used the z = 0 proton affinity and pKa,1 values determined from model compound measurements reported in references (62) and (63), respectively.

Per-atom partial charges for pSer and pThr were calculated using the LigParGen web server (64) and were normalized (in units of *e-*) such that the net charge for the whole residue was equal to the appropriate net charge (z = -2 or -1). To represent the resonance structures of the phosphoryl group, the z = -2 partial charges on all three oxygen atoms are equivalent (**Figure 1B**). The same is true for the partial charges of the phosphoryl group atoms for z = -1, except for the hydroxyl group oxygen, in which case the partial charge is slightly less negative because of the attached proton (see **Table S1**).

Ultimately, however, the parameters that enabled robust sampling of conformational space are those that combined z = -1 with a FOS corresponding to a deprotonated (z = -2) phosphoryl group. Use of z = -2 phospho-residues led to overstabilization of electrostatic interactions (i.e., Arg - pSer or Na^+^ - pSer), which ultimately yielded ‘trapped’ salt bridges and poorly sampled trajectories (**Figure S1A-D**). Recent work has implicated overly strong electrostatic interactions in fixed-charge force fields as contributing to sampling challenges and over-compaction in disordered proteins (31). As such, we propose that the reduction of charge density in our parameters is a necessary solution to avoid (likely) artifactual chelation between phosphate groups and positively charged residues or cations.

Unless otherwise stated, the simulations discussed herein reflect the “-2” state, that is, a phosphoryl harboring a -1 charge with the effective size and solvation energy of a -2 phosphoryl group. The resulting parameter files contain charge state-specific values and atom assignments; therefore, users must pass the parameters for the desired simulation charge state (z = -1 or -2). At present, we are unable to set different charge states for different phospho-residues in a protein sequence. Additional information about the implementation of these parameters and practical limitations may be found in the GitHub documentation.

### Comparison of OPLS-AA and CHARMM pSer/pThr tripeptide simulations

To evaluate the residue-level behavior of pSer and pThr in CAMPARI simulations, we simulated tripeptide sequences of Gly-Xaa-Gly with N- and C-terminal caps (acetyl and amide, respectively), where Xaa is Sep (pSer) or Tpo (pThr) in either z = -1 or z = -2 charge state. Five replicate simulations of 50,000,000 steps each were performed at 300 K and 0.05 M NaCl from different random starting conformations. After discarding equilibration frames, the merged trajectory contained 11,785 frames. Simulations were analyzed in Python using MDTraj (65).

### Model phospho-IDR simulations

All-atom simulations were performed using the ABSINTH implicit solvation model and CAMPARI Monte Carlo (MC) simulation engine (https://campari.sourceforge.net/). The simulations were conducted using the ion parameters derived by Mao et. al. (66) and the phospho-residue parameters described in this work. Each protein sequence (**Table 1**) was simulated using the same parameter file (*abs3.5_opls_phos-2.prm*) using the salt concentration (NaCl, modeled explicitly) that most closely aligns with that of the relevant experimental datasets. All protein sequences were simulated with N- and C-terminal capping groups to prevent aberrant cyclization. The simulations were analyzed in Python using SOURSOP and MDTraj (65, 67) and molecular graphics were generated in VMD or UCSF ChimeraX (68, 69). The simulation details for each of the different IDRs/peptides are provided in **Table S2**.

With the exception of the NHE1 transient helix 4 (TH4) peptide, all protein simulation replicates were initiated from a random coil starting structure. For the TH4 peptide, we performed five replicates of each random coil and helical starting conformations. Figure 2B was generated from the pool of all ten replicates; Figure 2C depicts trajectories from each starting state separately for sampling evaluation; and Figures 2D-E were generated using only the random coil-start trajectories to emphasize the impact of phosphorylation on induction of the helix.

**Figure 2:**
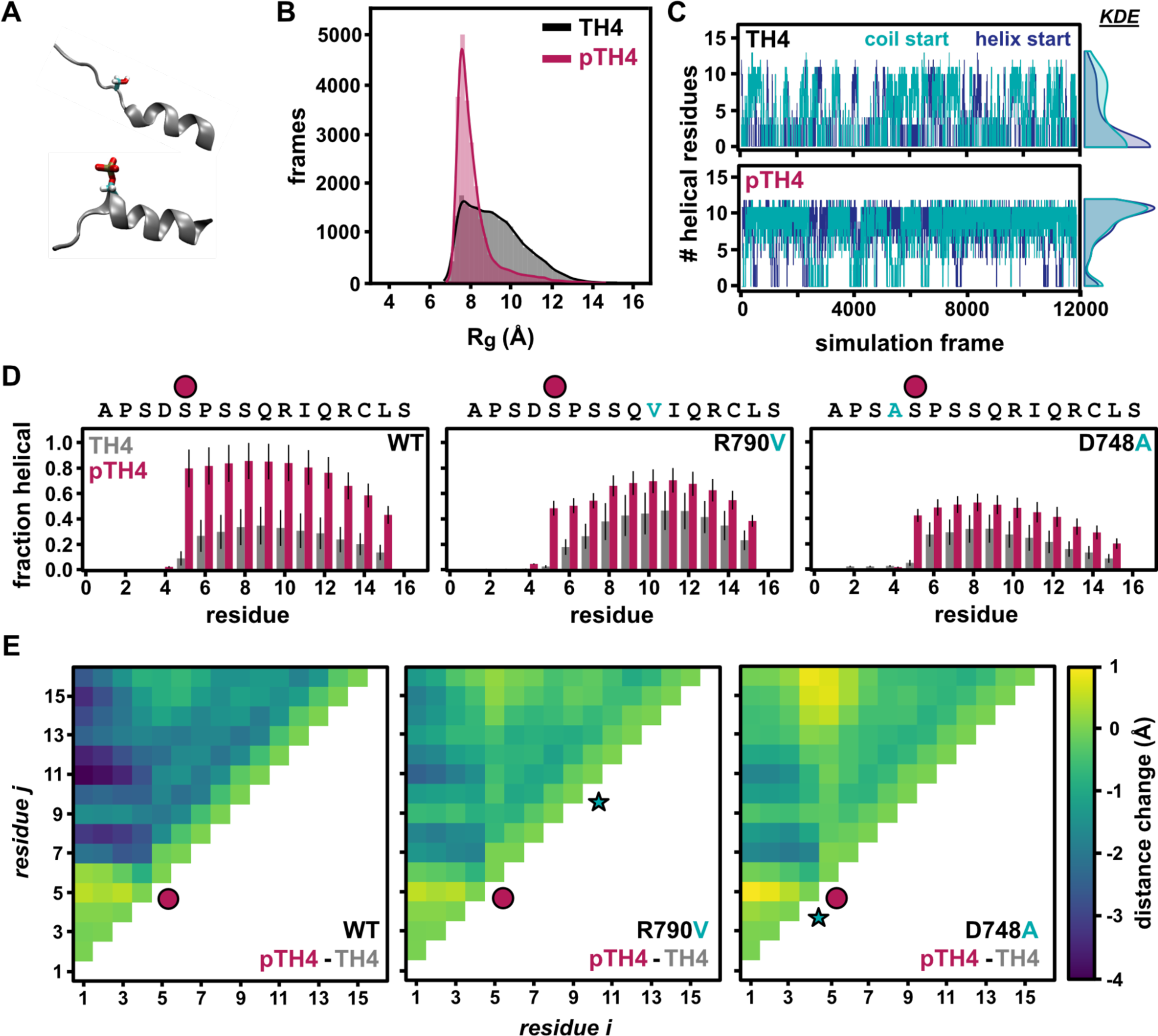
Ensemble properties of the TH4 peptide upon phosphorylation. **(A)** Representative snapshots of helical frames from unmodified (top) and phosphorylated (bottom) TH4. **(B)** Distributions of radius of gyration for the unmodified peptide (gray) and phosphorylated peptide (magenta). **(C)** Number of residues assigned as helical in the TH4 or pTH4 per frame of the simulation. Trajectories for random coil (‘coil start’) and helical (‘helix start’) starting conformations are plotted individually to demonstrate adequate sampling regardless of starting state structure. The distributions of the number of helical residues over the simulations are summarized as Kernel Density Estimate (KDE) plots on the right. **(D)** Per-residue helical content determined by DSSP for the WT peptide (left) and two point mutants (center and right). The magenta circle denotes the site of phosphorylation and, for the mutant peptides, the cyan residues are the sites of substitution. Error bars represent the standard deviation from the mean values determined from each of the replicates. **(E)** Difference heat maps demonstrating changes to intramolecular distances upon phosphorylation for TH4 WT and mutants. The distance change was calculated by subtracting the distance map of unmodified TH4 from that of phosphorylated TH4. Hence, a negative distance change denotes residue pairs that get closer together upon phosphorylation.

### Calculation of persistence lengths from simulations

Persistence lengths for Tau or pTau peptides were approximated from trajectories in two different ways: 1) using the end-to-end distance distribution and 2) from the decay of the angular correlation across the peptide. In method 1, persistence lengths were estimated from end-to-end distance distributions based on the following relation (43, 70):

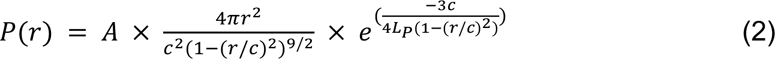

Where A is a scaling constant for normalization, r is the end-to-end distance, c is the contour length, and LP is persistence length in distance units (i.e., Å) In method 2, the angular correlation *g(s)* was calculated for all replicate trajectories using SOURSOP and plotted against interresidue separation (s). The persistence length (LP) was determined by fitting these data to an exponential decay model (71):

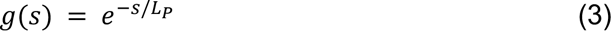

### Code and distribution

Phosphoparameters are provided in the abs3.5_opls_phos-2.prm parameter file, along with a short tutorial describing how to set up simulations using the pSer and pThr parameters in ABSINTH. The data and code that were used to generate the figures in this paper may be found here: https://github.com/holehouse-lab/supportingdata/tree/master/2024/Usher_phos_2024.

## RESULTS

### All-atom MC simulations of phosphorylated IDRs in CAMPARI

In ABSINTH, atoms are grouped together into small molecule-sized fragments (solvation groups), and each solvation group is assigned a free energy of solvation (ΔGsolv) that describes its interaction with the solvent. For charged side chains, this value is on the order of -100 kcal/mol, whereas the polar side chain of serine has ΔGsolv = -5 kcal/mol (50). The phosphoryl group arising from serine or threonine phosphorylation can harbor -1 or -2 charge at physiological pH owing to pKa values between 5.9 - 6.3 (61, 72) (**Figure 1B**). The charge states of all ionizable residues, including the phosphoryl groups, are fixed during the simulation. Unless otherwise specified, the simulations reported herein use a phosphoryl group with a -1 charge, with ΔGsolv on the order of ∼ -300 kcal/mol (**Figure 1B**).

The ABSINTH implicit solvent model comes in several “flavors”, with atomic partial charges associated with OPLS-AA and CHARMM36 being the most commonly used versions (abs3.5_opls.prm or abs3.5_charmm36.prm, respectively). These combine the originally reported ABSINTH parameters with the ion parameters derived by Mao *et. al* (66). The OPLS-AA version of ABSINTH has been applied to many IDR systems with excellent agreement with experimental ensemble descriptions (44, 52–58). We similarly observed good agreement between our simulated ensembles described herein and those described by published experiments and simulations. For comparison between the two force fields, we simulated tripeptides of the sequence Gly-Xaa-Gly, in which the central residue was phosphoserine or phosphothreonine with either -1 or -2 charge (**Table S1**). A comparison of the φ/ψ space for the central phosphorylated residue shows very good agreement of the two force fields for both phospho-residues and charge states at the level of torsion angles (**Figure 1C**).

Interatomic distances between the phosphoserine amide nitrogen (Sep2-N) and the phosphoserine phosphorus (Sep2-P) (N-P) or Tpo2-N and Tpo2-P are well-sampled during the simulations (**Figure S2A-B**). While the interatomic distances are consistent between OPLS-AA and CHARMM36 simulations of the same peptide, there are modest differences between Ser versus Thr and between charge states. The addition of the phosphate sterically constrains both Ser and Thr, but this effect is more pronounced in pThr due to its branched side chain (73, 74). In line with the reported steric and hydrogen bonding preferences of pThr compared to pSer, N-P distances (a proxy for a side chain-main chain hydrogen bond) are systematically smaller for pThr than pSer regardless of force field or charge state (**Figure S2C**) (74). The agreement of the force fields with one another and the ability to capture the reported steric preferences of pThr and pSer together suggest that, at the residue level, our simulations robustly sample the relevant local conformational space.

### Local phosphorylation-induced changes to the conformational ensemble

We next simulated two short peptides for which phosphorylation reportedly alters local conformational preferences (**Table 1**). One such conformational change is α-helix stabilization as a consequence of phosphorylation at the helix N-terminus, arising mainly from newly satisfied hydrogen bonds via the backbone or side chains or from favorable electrostatic interactions with the phosphoryl group (41, 75–77). Depending on the location of the phospho-site within the helix and the broader sequence context, phosphorylation-induced helix destabilization is another possible outcome (78).

Using our pSer and pThr parameters in ABSINTH/OPLS-AA, we simulated a transiently helical fragment from the sodium-proton exchanger 1, herein referred to as “transient helix 4” (TH4) (**Figure 2A**). Prior biophysical characterization of this fragment revealed that phosphorylation at the N-terminus of TH4 reinforced the helix through electrostatic interaction of the phosphoryl group with Arg in the *i + 5* position (41). Consistent with local ordering upon phosphorylation, we see compaction of the ensemble in the phosphorylated peptide (pTH4) simulation compared to the unmodified sequence (**Figure 2B**). Importantly, the degree of helicity for TH4 and pTH4 does not depend on the simulation starting structure (see *Methods*) (**Figure 2C**). However, to illustrate the substantial impact of this phosphorylation event on local secondary structure, **Figures 2D and E** represent trajectories in which all replicates used the ‘coil’ (unstructured) starting state.

From simulations of wild-type (WT) TH4 and pTH4, we determined the per-residue average helical fraction using DSSP (79) (**Figure 2D**). Indeed, we observe helix stabilization in pTH4 compared to TH4. This manifests at the residue level by an increase in helical character from about 10% to 80% at the site of phosphorylation (phosphoserine is marked by a magenta circle). Further, the per-residue helical character in the rest of the peptide increases upon N-terminal phosphorylation (**Figure 2D, left**). We also simulated the two point mutants described in (41) to deconvolute the contributions to phosphorylation-induced helix stability.

Our simulations showed that phosphorylation-induced helix stabilization was less pronounced than WT for both mutant peptides (**Figure 2D**). Published results implicate an emergent salt bridge between pS785 and R790 (*i + 5*) as the main helix stabilizer (41); in contrast, we find that these side chains do not fall on the same helix face. Instead, our phospho-WT simulations place the upstream D784 side chain in proximity to the R790 guanidinium group (**Figure S3A-C**). This result is in line with the failure by either point mutant to stabilize the helix to the extent of the WT sequence. In all three cases, however, we observed an increase in helical stability upon phosphorylation compared to the unmodified peptides, which we attribute to the interaction between the phosphoryl group and the backbone at residue S788 (*i + 3*) (**Figure S3D-F**).

As another test of our parameters to capture local structural changes upon phosphorylation, we used a peptide derived from a proline-rich region of Tau (residues 173-183 of Uniprot ID: P10636-8 (43)) that contains two phosphorylatable threonine (“TauT”) or serine (“TauS”) residues. The original study tested the hypothesis that phosphorylation within proline-rich sequences stabilizes poly-proline II structure and, therefore, increases the end-to-end distance of the peptide (43). We simulated the sequences in **Figure 3A** with or without two phosphates (or Glu phosphomimetics, “eTau”).

**Figure 3:**
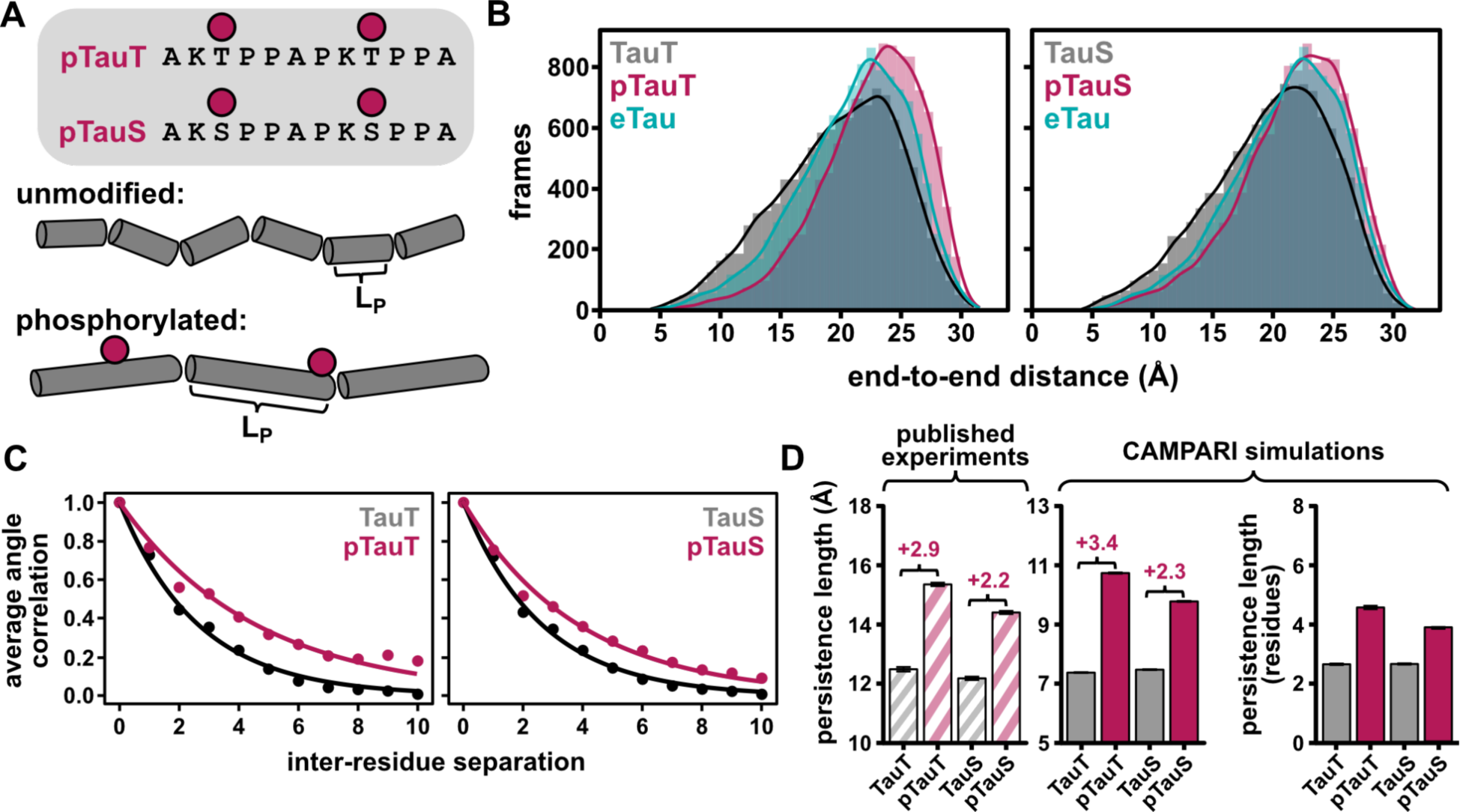
Changes to a proline-rich Tau peptide stiffness upon 2X-phosphorylation. **(A)** Top: sequences of Tau peptides used for simulations, where magenta circles show the locations of the phosphosites. Bottom: diagram of the reported effects of phosphorylation on persistence length (L_P_). **(B)** End-to-end distance distributions for unmodified (gray), phosphorylated (magenta), or Glu (phosphomimetic) (cyan) Tau peptides. **(C)** Plots of average angular correlation versus inter-residue separation used to calculate persistence length for Tau (black) and pTau (magenta). Exponential decay fits are shown as solid lines. **(D)** Summary of persistence length measurements from published experiments (43) and the simulations in this study.

We found the end-to-end distance distributions from our simulations trend qualitatively with the published finding that end-to-end distance increases upon peptide phosphorylation (43) (**Figure 3B**). The average end-to-end distances for unphosphorylated TauS and TauT are the same within error (⟨Ree⟩ = 19.8 ± 0.1 Å). The double phosphomimetic substituted peptide (eTau) is modestly more extended than TauS and TauT, with ⟨Ree⟩= 21 ± 0.2 Å. The phosphorylated peptides (pTauS and pTauT) are both more extended than the unphosphorylated and phosphomimetic peptides (⟨Ree⟩ = 21.7 ± 0.2 Å and 22.5 ± 0.1 Å, respectively). The published pTau study used experimental end-to-end distance distributions to approximate persistence length (LP), which quantifies the relative stiffness of a polymer and is defined as the distance along a chain over which bond angle correlation persists (43, 70).

We fit the distributions from our simulations in **Figure 3B** with **Equation 2**, which relates the end-to-end distance distribution to the contour length and persistence length, to quantify the change in LP with the addition of two phosphates (**Figures 3B,D and S4A-E, Table S2**). Unmodified TauS and TauT have nearly identical LP values (43) (**Figure 3D, Table S2**). In comparison to the published values, which determined end-to-end distance distributions from FRET, the persistence lengths we observe are systematically smaller because our simulations lack the terminal donor and acceptor fluorophores (**Table S2**). Nevertheless, the increases in LP between TauT or TauS and pTauT or pTauS from simulations are in excellent agreement with experiments (**Figure 3D**). The TauS LP increased by 2.2 and 2.3 Å in experiments and simulations, respectively, after phosphorylation. The TauT LP increased with phosphorylation by 2.9 (published experiments) and 3.4 (simulations) Å (43). Notably, the LP for phosphomimetic Tau was greater than the unmodified peptide (+1.6 Å), but the increase was not as dramatic as for the true phosphorylated peptides.

From atomistic simulations, we can extract certain structural parameters and features of an ensemble that are inaccessible in experiments. We separately calculated persistence lengths for TauS, TauT, pTauS, and pTauT peptides using the decay of angular correlations over the length of the peptide chain. The average angular correlation is the average angle between two bond vectors (C > N) between two residues in a chain. We used the decay of the angular correlation to determine LP using the relation in **Equation 3** (**Figure 3C**), based on the approximation that the cosine of the angle between the bond vectors will decay exponentially over the length of the peptide (71). Indeed, persistence lengths calculated in this way (in units of residues) show the same trend as those calculated from the end-to-end distance distributions (**Figure 3D**).

Taken together, our analysis of short disordered peptides suggests that our phosphoparameters reasonably capture local steric and conformational biases. Importantly, we observe local chain compaction via induced helicity and chain expansion via a combination of favorable solvation and electrostatic repulsion. These results suggest the energetics of the phosphate moieties are well-balanced, enabling subtle context-dependent effects.

### Effects of multi-site phosphorylation on IDR ensemble dimensions

We next assessed the ability of our pSer and pThr parameters to capture phosphorylation-induced ensemble effects on two longer IDR sequences that undergo multi-site phosphorylation. We selected IDRs that undergo multiple phosphorylation events and have been characterized using *in vitro* and *in silico* methods in their unphosphorylated and phosphorylated states. An IDR of the yeast transcription factor Ash1 (residues 420-500) is 10X-phosphorylated by a cyclin-dependent kinase, but does not change appreciably in global ensemble dimensions (44) (**Figures 4A,C**). In contrast, the ensemble of PAGE4, a human regulatory protein, expands dramatically with the installation of eight phosphates by the kinase CLK2 (46, 47) (**Figures 4B,D**). These distinct biophysical responses to comparable changes in phosphorylation state offer a good test for the validity of our phosphoparameters.

**Figure 4:**
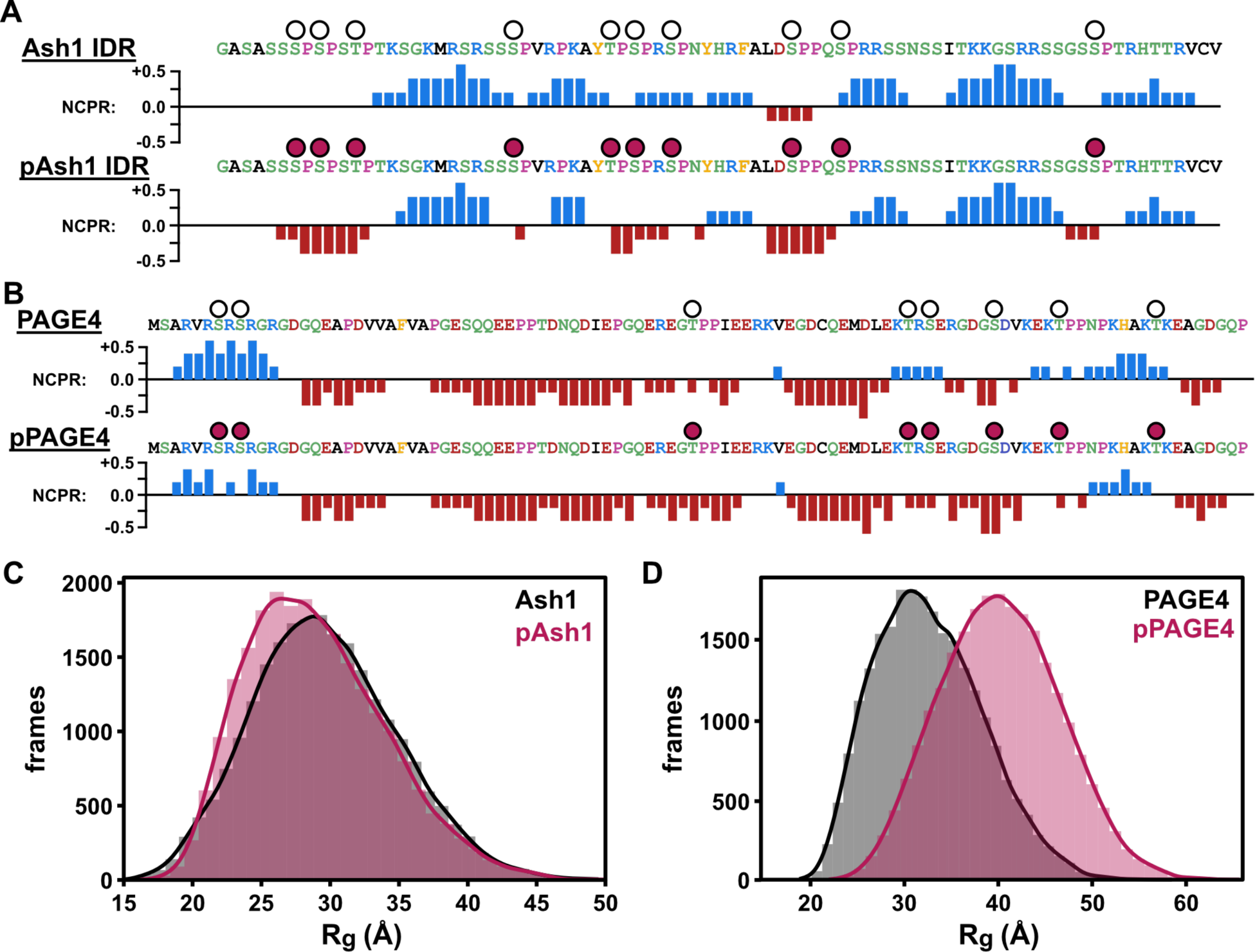
Sequence and ensemble properties of Ash1 and PAGE4 with and without multi-site phosphorylation. **(A)** Sequences and net charge per residue (NCPR) profiles for Ash1 and pAsh1. **(B)** Sequences and NCPR profiles for PAGE4 and pPAGE4. For **(A)** and **(B),** circles above the sequences denote sites of phosphorylation; filled circles denote that the site is phosphorylated in the simulation. NCPR was calculated for the pAsh1 and pPAGE4 by replacing each phospho-site in the sequence with a phosphomimetic residue (Asp) to reflect a -1 charge. **(C)** Distribution of Rg values for Ash1 (gray) or pAsh1 (magenta) simulations. **(D)** Distribution of Rg values for PAGE4 (gray) or pPAGE4 (magenta) simulations.

For both Ash1 and PAGE4, the sequence and sites of phosphorylation are shown above a plot of net charge per residue (NCPR) to visually represent the changes to the charge landscape associated with phosphorylation (**Figures 4A-B**). In line with published effects of phosphorylation for Ash1, the ensemble dimensions are the same within error for the unphosphorylated (⟨Rg⟩ = 29.6 ± 0.4 Å) and phosphorylated states (⟨Rg⟩ = 29.0 ± 0.6 Å). Despite a nearly 50% decrease in NCPR for phosphorylated Ash (pAsh1), possible charge-driven changes to the ensemble dimensions are offset due to the patterning of proline residues across the sequence and local, compensatory phosphorylation-induced features (44) (**Figures 4C & S5A**).

On the other hand, unmodified PAGE4 has a small, slightly negative NCPR that becomes more negative with 8X-phosphorylation (**Figure 4B**). Unlike Ash1, the global dimensions of PAGE4 are highly sensitive to hyperphosphorylation, characterized by an increase from ⟨Rg⟩ = 32.6 ± 0.9 Å (PAGE4) to ⟨Rg⟩ = 40.2 ± 1.3 Å (pPAGE4). The magnitude of this change is in quantitative agreement with dimensions extracted from FRET, SAXS, and molecular dynamics (MD) simulations (46–48). Across the experimental and computational datasets for PAGE4, the dimensions measured for pPAGE4 by SAXS are markedly higher than from FRET, MD, and our simulations. Nevertheless, experiments and simulations alike point toward a dramatic (> 20%) increase in the PAGE4 ⟨Rg⟩ that accompanies 8X-phosphorylation (**Figures 4D & S5B**).

### Changes to IDR charge landscape upon multiple phosphorylation events

We next explored the conformational impact of specific patterns of phosphorylation within Ash1 and PAGE4. Experimental studies of phospho-IDRs are generally limited by the ability to enzymatically generate phosphorylated proteins in sufficient quantities and homogeneity for biophysical characterization. In simulations, however, because we have control of the input sequence, we may specify exactly which residues are phosphorylated. We surveyed the Ash1 and PAGE4 phospho-sequence space (**Figures S5C-D**) to find combinations of phosphosites that alter the NCPR and then simulated these sequences using our phospho-residue parameters (**Figure 5A-B**, **Tables S3-4**).

**Figure 5:**
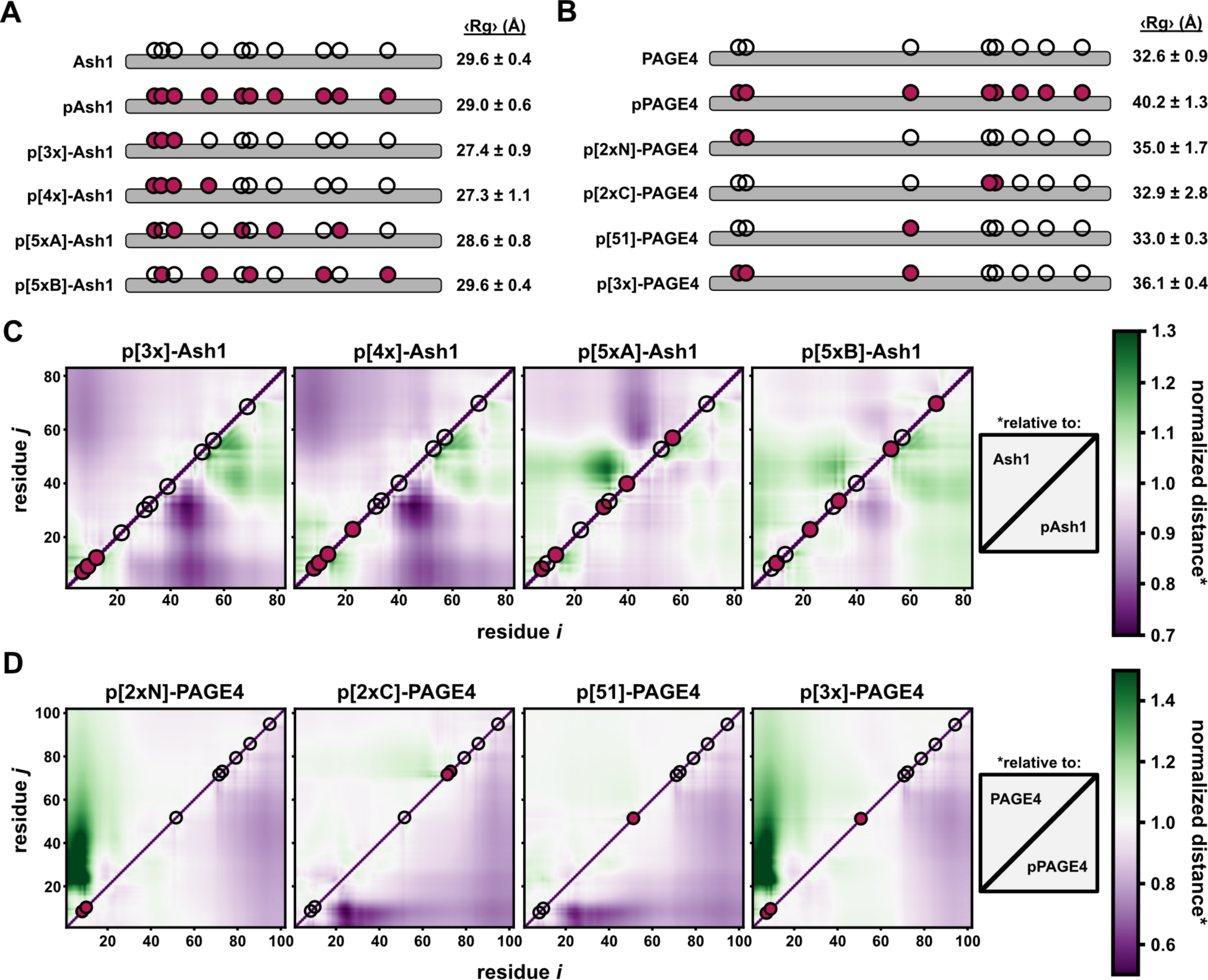
Effects of phosphorylation patterning on IDR ensemble properties. **(A)** Phospho-forms of Ash1 and their ensemble dimensions (R_G_). Reported error is the standard deviation from the mean R_G_ value from all ten replicates. Filled magenta circles denote phosphorylated residues for each phospho-form. **(B)** Phospho-forms of PAGE4 and their ensemble dimensions (R_G_). Reported error is the standard deviation from the mean R_G_ value from all ten replicates. Filled magenta circles denote phosphorylated residues for each phospho-form. **(C)** and **(D)** Heat maps from simulations of Ash1 **(C)** and PAGE4 **(D)** phosphorylation combinations. The upper left triangle of the heat map was generated by dividing the distance map of a given phospho-form by that of the unmodified protein reference state; the lower right triangle was generated similarly, but was instead normalized against the fully phosphorylated distance map. A normalized distance equal to 1 would suggest there is no change from the reference state (i.e., relative to the unmodified or hyperphosphorylated sequence). Values >1 and <1 reflect regions that are more expanded and more compact, respectively, relative to the reference state.

Drawing from the sequence similarities (i.e., high proline content) between the short Tau peptides, which expand modestly upon phosphorylation (43), and segments of pAsh1, we constructed normalized distance maps for each simulated phospho-state to observe potential local effects of particular phosphorylation events (**Figure 5C**). For each sequence, the upper triangle of the distance map is normalized to the completely unmodified Ash1, while the lower triangle of the same panel is normalized to the hyperphosphorylated sequence (pAsh1). This presentation allows us to see regions of relative expansion and compaction resulting from specific combinations of phosphosites. We plotted the same for select combinations of phosphosites in PAGE4 to probe the effects of fewer phosphates on the local and global ensembles (**Figure 5D**).

The distance map for pAsh1 normalized to unmodified Ash1 highlights published conclusions from simulations of phosphomimetic Ash1 that local expansion (green) is offset by local compaction elsewhere (purple) that ultimately yields no net change in ⟨Rg⟩ (**Figure S5A**) (44). p[3x]-Ash1 and p[4x]-Ash1, which cluster the phosphates at the N-terminus, are modestly more compact than either Ash1 or pAsh1 based on ⟨Rg⟩ (**Figure 5A**). From their distance maps, the region that harbors these phosphosites is more expanded than the unmodified Ash1, whereas the rest of the protein is more compact than unmodified Ash1 (**Figure 5C**). Distance maps for both 5X-phosphorylated Ash1 constructs show distinct regions of relative expansion and relative compaction, which supports published conclusions that local expansion and compaction associated with phosphorylation collectively offset any changes to the global dimensions (44). For Ash1, which is net positively charged in the unmodified form, the change in NCPR from phosphorylation does not appear to strongly correlate with Rg (**Figure S5E**) as predicted by an electrostatics-driven model (36).

Unmodified PAGE4 harbors a net negative charge of -8 and has a slightly negative NCPR that becomes more negative with more phosphorylation events (**Table S4**). p[2xN]-PAGE4 contains pS7 and pS9, which fall within a basic RS-rich motif directly N-terminal to an acidic patch. In unmodified PAGE4, the N-terminus is somewhat compact due to interactions between the basic and acidic regions. Phosphorylation within the basic region leads to decompaction of this region of PAGE4 and contributes to a more expanded global ensemble (**Figure 5D**). A pair of phosphosites (pT71 and pS73) toward the C-terminus contribute to increased negative charge density but do not themselves contribute to the global dimensions of PAGE4.

pT51 is one of two reported sites that are phosphorylated by HIPK1 to modulate PAGE4 interactions with stress response transcription factors (80). p[51]-PAGE4 has approximately the same global dimensions as unmodified PAGE4, which may speak to the use of this phosphosite to modify a particular binding motif. When combined with pS7 and pS9, the degree of expansion (from 32.6 Å to 36.1 Å) is still less pronounced than 8X-phosphorylated PAGE4 (to 40.2 Å) (**Figures 5B & S5E, Table S4**). The working model is that PAGE4 hyperphosphorylation (all eight sites) diminishes its interactions with stress response factors and leads to its degradation (46). Thus, the preference for a somewhat collapsed conformation in hypo-phosphorylated states may reflect regulatory protection of PAGE4 in the absence of a degradation-promoting kinase signal.

## DISCUSSION

In this study, we implemented new parameters for phosphothreonine and phosphoserine for use in CAMPARI Monte Carlo simulations of disordered proteins and assessed the results of such simulations compared with published findings. We find that these parameters qualitatively reproduce experimental and computational results for phosphorylation-induced α-helix stabilization (TH4) and increased chain stiffness (Tau) in short peptides (41, 43). We also tested the ability of our parameters to capture reported changes to the ensembles of two longer IDRs (80 and 102 residues) with multiple phosphorylation events; indeed, our phospho-IDR simulations reproduce published changes in ensemble dimensions with near-quantitative accuracy (44, 46, 48).

### On the use of phosphomimetic mutations

In this study of the effects of phosphorylation on select peptides and IDRs, we see excellent agreement of our simulations with findings from published experiments and simulations. In all four test cases described here, we also tested the extent to which phosphomimetic mutations (S/T > E) could recover the effects of true phosphorylation (S/T/ > pS/pT). In the TH4 peptide, for which phosphorylation stabilizes α-helical structure, substituting a phosphomimetic residue (eTH4) suggests that E is more stabilizing than S, but not to the extent of the stability gained by a true phosphoserine (**Figure S4F**). For phosphomimetic Tau (eTau), the increase in persistence length compared to the unmodified peptide was *less pronounced* than the increase associated with true phosphothreonine or phosphoserine (**Table S2**). This result is consistent with previous studies of the Tau proline-rich region, which found that the branched Thr side chain contributes to the phosphorylation-induced stiffening by steric effects (81).

In contrast, the ensemble features for the two longer phospho-IDRs we tested are well-described by multi-site phosphomimetic mutations. This was demonstrated for Ash1 in the original publication, which used phosphomimetic Ash1 (“eAsh1”) in simulations prior to the availability of reliable phospho-residue parameters (44). eAsh1 and pAsh1 ensemble dimensions are highly similar between simulations and SAXS experiments. The agreement between eAsh1 and pAsh1 also holds true for relative intramolecular distances and for changes in local radius of gyration for two sub-regions of Ash1 (**Figure S5F**). For PAGE4, our simulations suggest that global ensembles ePAGE4 and pPAGE4 are indistinguishable, which is consistent with a largely charge-driven expansion upon hyperphosphorylation. Indeed, the distance maps for ePAGE4 or pPAGE4 relative to unmodified PAGE4 suggest that the intramolecular changes with phosphomimetics versus true phosphorylation are generally the same (**Figure S5B**).

The use of phosphomimetic mutations is a common workaround for studying phosphoproteins *in vitro* and *in vivo*, but the success of this strategy at recapitulating true phosphorylation is highly system-dependent. For the yeast cell cycle regulator Whi5, mutation of four phosphosites to glutamates (representing a constitutively inactive phosphorylated state) yields a phenotype that is similar to that of a *whi5* deletion (82). Similarly, changes in phase separation propensity upon multi-site phosphorylation in IDRs such as FUS, TDP-43, and Ki-67 are effectively reproduced by multi-site phosphomimetics (83–85).

However, as shown herein and elsewhere, phosphorylation events that modulate local structural propensity and/or a particular interaction motif may be poorly represented by a phosphomimetic mutation. In the protein 4E-BP2, the phosphorylation of two threonine residues drives local folding into a four-stranded β sheet domain, which is, notably, not observed with phosphomimetics (14). In the context of α-Synuclein self-assembly, pSer129 inhibits fibril formation, whereas S129D and S129E do not (86). Increased end-to-end distance of a Tau peptide upon phosphorylation is only partially recovered when using a phosphomimetic mutation (81). Phosphorylation of the kinase RSK within its activation loop is required for catalytic function, but the phosphomimetic mutant is insufficient to stabilize the local structure (87). For the 14-3-3 protein, the dimer-monomer equilibrium is controlled via phosphorylation of S58, and S58E fails to shift this equilibrium (88).

Although the simulations herein do not explore protonation/deprotonation events, the near-physiological pKa of the phosphoryl group poises it to respond to changes to the local pH environment in a manner that is impossible for aspartate or glutamate (89). Indeed, depending on sequence context, multisite phosphorylation in an IDR can confer conformational sensitivity to pH, which is inherently linked to the ability of the phosphoryl group to undergo protonation/deprotonation (90, 91). Setting aside structural differences in the aspartate/glutamate and phosphoserine/phosphothreonine side chains, the potential for greater negative charge density on true phospho-residues further sets them functionally apart from the phosphomimetic residues.

For these reasons, we advise caution when using phosphomimetic substitutions to probe the effects of phosphorylation–for disordered and folded proteins alike. In our four test proteins, the long, multi-phosphorylated IDRs with global ensemble changes seem to be described well by phosphomimetics; in the shorter peptides, we find that the phosphomimetics do not confer the same local structural consequences as true phosphorylation. Importantly, while we provide examples of other systems that do and do not suit the use of phosphomimetics, there certainly are exceptions to these broad classifications.

### Limitations and caveats

As is always the case for computational methods, it behooves all involved to be aware of the layers of simplifying assumptions being made, which may impact the validity of simulation-derived conclusions. It is important to acknowledge some limitations associated with the use of a continuum (implicit) solvent model, particularly for charged molecules that have significant solvation requirements (i.e., phosphoserine and phosphothreonine). Although we account for solvation free energy for different chemical groups, the nature of the solvent model assigns the same solvation volume to different types of molecules; however, it is known that identity of the solvated molecule can subtly or dramatically affect the shape, size, and organization of the water shell (37, 59). Despite this assumption, it is worth noting that recent work has shown ABSINTH to perform relatively well in the context of *in-silico* small-molecule binding (92). As such, while we might expect the impact of a consistent solvent volume to be more pronounced for highly charged chemical moieties (e.g., phosphorylated residues), extant work suggests it is a reasonable approximation for a wide variety of chemical interactions.

A second key caveat here is the absence of charge regulation as taken into account in these simulations (93). Charge regulation reflects that if many charged residues of the same sign are locally constrained near one another, the intrinsic pKa of those side chains may be altered (93–95). Specifically, pKa values may be upshifted (if acidic) or downshifted (if basic), enabling neutralization via protonation (if acidic) or deprotonation (if basic) under bulk pH conditions where changes in ionization may not naively be expected. Future work focusing on multisite phosphorylation of negatively charged regions should take this into account.

A third caveat reflects the more general caveat that fixed charge force fields may have limitations in their ability to fully capture the complex polarization effects seen in chemical groups with multiple resonance structures. The ability of fixed-charged models to correctly describe phosphate-arginine interaction – in particular – remains unclear, so extra care in interpretation should be taken if specific phosphate-arginine interactions are expected. Notably, we have not described parameters for simulating phosphotyrosine (pTyr). The rationale is twofold: first, we anticipate correctly capturing the consequences of multiple resonance structures to be especially challenging; second, compared to pSer and pThr there are very few biophysical studies of pTyr in IDRs with which to compare our simulations. Combined, we felt addressing an inherently challenging problem with a paucity of data would be a fraught exercise.

Finally, in developing these parameters, when using the –2 charge state, we frequently observed tight binding of sodium ions (Na^+^) whereby the oxygen atoms associated with the phosphate group and that of the backbone carbonyl provide an oxygen “cage” for Na^+^ chelation (**Figure S1D**). In our simulations, chelation manifests as an irreversible binding of Na^+^ to phosphate groups, leading to energetically trapped states whereby local conformational rearrangement is dramatically reduced.

While we cannot formally exclude the possibility that such chelation occurs, we do not see support for such robust chelation from experiments. In prior work, pAsh1 was measured by SAXS between 50 and 1000 mM NaCl, yet no appreciable change in global dimensions, solubility, or the shape of the scattering curve was observed (44). More recently, titrations of NaCl with the disordered protein DDX4 estimated NaCl binding to clusters of acidic residues to be extremely weak, with a Kd over 1000 mM (where calcium ions have an inferred average Kd of ∼150 mM), suggesting Na^+^ does not readily bind to polypeptides, even around regions of local negative charge density (96). Finally, molecular dynamics simulations of phosphorylated NCBD – in good agreement with single-molecule FRET experiments – find no evidence for Na^+^ chelation of phosphate groups (17). As such, while we cannot formally exclude the possibility, we tentatively suggest that Na^+^ chelation may be an artifactual consequence of fixed-charge electrostatics. At the very least, in our simulations, the consequences of chelation on local conformational rearrangements are almost certainly artifactual. While our choice of z = -1 for pSer and pThr mitigates this issue here, we caution others to be skeptical of such a result in their own systems and seek out strong experimental support for such a result if it is to be pursued as a *bona fide* observation.

## AUTHOR CONTRIBUTIONS

E.T.U. designed research, developed code, performed analyses, and wrote the manuscript. M.J.F. designed research, developed code, and contributed to manuscript revisions. A.S.H. designed research, supervised this work, and wrote the manuscript.

## ACKNOWLEDGEMENTS

We thank Rohit Pappu and members of the Pappu lab for many useful conservations and advice. We also thank the members of the Holehouse lab for helpful discussions and feedback on the manuscript. E.T.U. was supported on this work by a Keck Postdoctoral Fellowship. A.S.H. was supported by the NIH National Institute on Allergic and Infectious Diseases (NIAID) with R01AI163142.

## REFERENCES

1. Regmi, R., S. Srinivasan, A.P. Latham, V. Kukshal, W. Cui, B. Zhang, R. Bose, and G.S. Schlau-Cohen. 2020. Phosphorylation-Dependent Conformations of the Disordered Carboxyl-Terminus Domain in the Epidermal Growth Factor Receptor. J. Phys. Chem. Lett. 11:10037–10044, doi: 10.1021/acs.jpclett.0c02327.

2. Kõivomägi, M., M.P. Swaffer, J.J. Turner, G. Marinov, and J.M. Skotheim. 2021. G1 cyclin-Cdk promotes cell cycle entry through localized phosphorylation of RNA polymerase II. Science. 374.

3. Zhang, L., S. Winkler, F.P. Schlottmann, O. Kohlbacher, J.E. Elias, J.M. Skotheim, and J.C. Ewald. 2019. Multiple Layers of Phospho-Regulation Coordinate Metabolism and the Cell Cycle in Budding Yeast. Front Cell Dev Biol. 7:338, doi: 10.3389/fcell.2019.00338.

4. Topacio, B.R., E. Zatulovskiy, S. Cristea, S. Xie, C.S. Tambo, S.M. Rubin, J. Sage, M. Koivomagi, and J.M. Skotheim. 2019. Cyclin D-Cdk4,6 Drives Cell-Cycle Progression via the Retinoblastoma Protein’s C-Terminal Helix. Mol. Cell. 74:758–770 e4, doi: 10.1016/j.molcel.2019.03.020.

5. Ardito, F., M. Giuliani, D. Perrone, G. Troiano, and L. Lo Muzio. 2017. The crucial role of protein phosphorylation in cell signaling and its use as targeted therapy. Int. J. Mol. Med. 40:271–280, doi: 10.3892/ijmm.2017.3036.

6. Harlen, K.M., K.L. Trotta, E.E. Smith, M.M. Mosaheb, S.M. Fuchs, and L.S. Churchman. 2016. Comprehensive RNA Polymerase II Interactomes Reveal Distinct and Varied Roles for Each Phospho-CTD Residue. Cell Rep. 15:2147–2158, doi: 10.1016/j.celrep.2016.05.010.

7. Bondos, S.E., A.K. Dunker, and V.N. Uversky. 2021. On the roles of intrinsically disordered proteins and regions in cell communication and signaling. Cell Commun. Signal. 19:88, doi: 10.1186/s12964-021-00774-3.

8. Iakoucheva, L.M., P. Radivojac, C.J. Brown, T.R. O’Connor, J.G. Sikes, Z. Obradovic, and A.K. Dunker. 2004. The importance of intrinsic disorder for protein phosphorylation. Nucleic Acids Res. 32:1037–1049, doi: 10.1093/nar/gkh253.

9. Cho, M.H., J.O. Wrabl, J. Taylor, and V.J. Hilser. 2020. Hidden dynamic signatures drive substrate selectivity in the disordered phosphoproteome. Proc. Natl. Acad. Sci. U. S. A. 117:23606–23616, doi: 10.1073/pnas.1921473117.

10. Ginell, G.M., A.J. Flynn, and A.S. Holehouse. 2023. SHEPHARD: a modular and extensible software architecture for analyzing and annotating large protein datasets. Bioinformatics. 39doi: 10.1093/bioinformatics/btad488.

11. Newcombe, E.A., E. Delaforge, R. Hartmann-Petersen, K. Skriver, and B.B. Kragelund. 2022. How phosphorylation impacts intrinsically disordered proteins and their function. Essays Biochem.doi: 10.1042/EBC20220060.

12. Liu, N., Y. Guo, S. Ning, and M. Duan. 2020. Phosphorylation regulates the binding of intrinsically disordered proteins via a flexible conformation selection mechanism. Communications Chemistry. 3doi: 10.1038/s42004-020-00370-5.

13. Betts, M.J., O. Wichmann, M. Utz, T. Andre, E. Petsalaki, P. Minguez, L. Parca, F.P. Roth, A.-C. Gavin, P. Bork, and R.B. Russell. 2017. Systematic identification of phosphorylation-mediated protein interaction switches. PLoS Comput. Biol. 13:e1005462, doi: 10.1371/journal.pcbi.1005462.

14. Bah, A., R.M. Vernon, Z. Siddiqui, M. Krzeminski, R. Muhandiram, C. Zhao, N. Sonenberg, L.E. Kay, and J.D. Forman-Kay. 2015. Folding of an intrinsically disordered protein by phosphorylation as a regulatory switch. Nature. 519:106–109, doi: 10.1038/nature13999.

15. Bah, A., and J.D. Forman-Kay. 2016. Modulation of Intrinsically Disordered Protein Function by Post-translational Modifications. J. Biol. Chem. 291:6696–6705, doi: 10.1074/jbc.R115.695056.

16. Valverde, J.M., G. Dubra, M. Phillips, A. Haider, C. Elena-Real, A. Fournet, E. Alghoul, D. Chahar, N. Andrés-Sanchez, M. Paloni, P. Bernadó, G. van Mierlo, M. Vermeulen, H. van den Toorn, A.J.R. Heck, A. Constantinou, A. Barducci, K. Ghosh, N. Sibille, P. Knipscheer, L. Krasinska, D. Fisher, and M. Altelaar. 2023. A cyclin-dependent kinase-mediated phosphorylation switch of disordered protein condensation. Nat. Commun. 14:6316, doi: 10.1038/s41467-023-42049-0.

17. Buholzer, K.J., J. McIvor, F. Zosel, C. Teppich, D. Nettels, D. Mercadante, and B. Schuler. 2022. Multilayered allosteric modulation of coupled folding and binding by phosphorylation, peptidyl-prolyl cis/trans isomerization, and diversity of interaction partners. J. Chem. Phys. 157:235102, doi: 10.1063/5.0128273.

18. Portz, B., F. Lu, E.B. Gibbs, J.E. Mayfield, M. Rachel Mehaffey, Y.J. Zhang, J.S. Brodbelt, S.A. Showalter, and D.S. Gilmour. 2017. Structural heterogeneity in the intrinsically disordered RNA polymerase II C-terminal domain. Nat. Commun. 8:15231, doi: 10.1038/ncomms15231.

19. Gibbs, E.B., and S.A. Showalter. 2015. Quantitative biophysical characterization of intrinsically disordered proteins. Biochemistry. 54:1314–1326, doi: 10.1021/bi501460a.

20. Kikhney, A.G., and D.I. Svergun. 2015. A practical guide to small angle X-ray scattering (SAXS) of flexible and intrinsically disordered proteins. FEBS Lett. 589:2570–2577, doi: 10.1016/j.febslet.2015.08.027.

21. Cubuk, J., M.D. Stuchell-Brereton, and A. Soranno. 2022. The biophysics of disordered proteins from the point of view of single-molecule fluorescence spectroscopy. Essays Biochem. 66:875–890, doi: 10.1042/EBC20220065.

22. Naudi-Fabra, S., M. Tengo, M.R. Jensen, M. Blackledge, and S. Milles. 2021. Quantitative Description of Intrinsically Disordered Proteins Using Single-Molecule FRET, NMR, and SAXS. J. Am. Chem. Soc. 143:20109–20121, doi: 10.1021/jacs.1c06264.

23. Gomes, G.W., M. Krzeminski, A. Namini, E.W. Martin, T. Mittag, T. Head-Gordon, J.D. Forman-Kay, and C.C. Gradinaru. 2020. Conformational Ensembles of an Intrinsically Disordered Protein Consistent with NMR, SAXS, and Single-Molecule FRET. J. Am. Chem. Soc. 142:15697–15710, doi: 10.1021/jacs.0c02088.

24. Cook, E.C., G.A. Usher, and S.A. Showalter. 2018. The Use of (13)C Direct-Detect NMR to Characterize Flexible and Disordered Proteins. Methods Enzymol. 611:81–100, doi: 10.1016/bs.mie.2018.08.025.

25. Mercadante, D., J.A. Wagner, I.V. Aramburu, E.A. Lemke, and F. Gräter. 2017. Sampling Long-versus Short-Range Interactions Defines the Ability of Force Fields To Reproduce the Dynamics of Intrinsically Disordered Proteins. J. Chem. Theory Comput. 13:3964–3974, doi: 10.1021/acs.jctc.7b00143.

26. Levine, Z.A., and J.-E. Shea. 2017. Simulations of disordered proteins and systems with conformational heterogeneity. Curr. Opin. Struct. Biol. 43:95–103, doi: 10.1016/j.sbi.2016.11.006.

27. Rauscher, S., V. Gapsys, M.J. Gajda, M. Zweckstetter, B.L. de Groot, and H. Grubmüller. 2015. Structural Ensembles of Intrinsically Disordered Proteins Depend Strongly on Force Field: A Comparison to Experiment. J. Chem. Theory Comput. 11:5513–5524, doi: 10.1021/acs.jctc.5b00736.

28. Choi, J.M., and R.V. Pappu. 2019. Improvements to the ABSINTH Force Field for Proteins Based on Experimentally Derived Amino Acid Specific Backbone Conformational Statistics. J. Chem. Theory Comput. 15:1367–1382, doi: 10.1021/acs.jctc.8b00573.

29. Adhikari, S., and J. Mondal. 2023. Machine Learning Subtle Conformational Change due to Phosphorylation in Intrinsically Disordered Proteins. J. Phys. Chem. B. 127:9433–9449, doi: 10.1021/acs.jpcb.3c05136.

30. Sieradzan, A.K., A. Korneev, A. Begun, K. Kachlishvili, H.A. Scheraga, A. Molochkov, P. Senet, A.J. Niemi, and G.G. Maisuradze. 2021. Investigation of Phosphorylation-Induced Folding of an Intrinsically Disordered Protein by Coarse-Grained Molecular Dynamics. J. Chem. Theory Comput. 17:3203–3220, doi: 10.1021/acs.jctc.1c00155.

31. Piana, S., P. Robustelli, D. Tan, S. Chen, and D.E. Shaw. 2020. Development of a Force Field for the Simulation of Single-Chain Proteins and Protein-Protein Complexes. J. Chem. Theory Comput. 16:2494–2507, doi: 10.1021/acs.jctc.9b00251.

32. Robustelli, P., S. Piana, and D.E. Shaw. 2018. Developing a molecular dynamics force field for both folded and disordered protein states. Proc. Natl. Acad. Sci. U. S. A. 115:E4758– E4766, doi: 10.1073/pnas.1800690115.

33. Mittal, A., R.K. Das, A. Vitalis, and R.V. Pappu. 2014. ABSINTH Implicit Solvation Model and Force Field Paradigm for Use in Simulations of Intrinsically Disordered Proteins. In: Fuxreiter M, editor. Computational Approaches to Protein Dynamics. CRC Press. pp. 181– 203, doi: 10.1201/b17979-13.

34. Rieloff, E., and M. Skepö. 2020. Phosphorylation of a Disordered Peptide-Structural Effects and Force Field Inconsistencies. J. Chem. Theory Comput. 16:1924–1935, doi: 10.1021/acs.jctc.9b01190.

35. Rieloff, E., and M. Skepö. 2021. Molecular Dynamics Simulations of Phosphorylated Intrinsically Disordered Proteins: A Force Field Comparison. Int. J. Mol. Sci. 22doi: 10.3390/ijms221810174.

36. Jin, F., and F. Grater. 2021. How multisite phosphorylation impacts the conformations of intrinsically disordered proteins. PLoS Comput. Biol. 17:e1008939, doi: 10.1371/journal.pcbi.1008939.

37. Wong, S.E., K. Bernacki, and M. Jacobson. 2005. Competition between intramolecular hydrogen bonds and solvation in phosphorylated peptides: simulations with explicit and implicit solvent. J. Phys. Chem. B. 109:5249–5258, doi: 10.1021/jp046333q.

38. Homeyer, N., A.H. Horn, H. Lanig, and H. Sticht. 2006. AMBER force-field parameters for phosphorylated amino acids in different protonation states: phosphoserine, phosphothreonine, phosphotyrosine, and phosphohistidine. J. Mol. Model. 12:281–289, doi: 10.1007/s00894-005-0028-4.

39. Song, G., B. Zhong, B. Zhang, A.U. Rehman, and H.-F. Chen. 2023. Phosphorylation Modification Force Field FB18CMAP Improving Conformation Sampling of Phosphoproteins. J. Chem. Inf. Model. 63:1602–1614, doi: 10.1021/acs.jcim.3c00112.

40. Stanley, N., S. Esteban-Martín, and G. De Fabritiis. 2014. Kinetic modulation of a disordered protein domain by phosphorylation. Nat. Commun. 5:5272, doi: 10.1038/ncomms6272.

41. Hendus-Altenburger, R., M. Lambrughi, T. Terkelsen, S.F. Pedersen, E. Papaleo, K. Lindorff-Larsen, and B.B. Kragelund. 2017. A phosphorylation-motif for tuneable helix stabilisation in intrinsically disordered proteins - Lessons from the sodium proton exchanger 1 (NHE1). Cell. Signal. 37:40–51, doi: 10.1016/j.cellsig.2017.05.015.

42. Schwalbe, M., H. Kadavath, J. Biernat, V. Ozenne, M. Blackledge, E. Mandelkow, and M. Zweckstetter. 2015. Structural Impact of Tau Phosphorylation at Threonine 231. Structure. 23:1448–1458, doi: 10.1016/j.str.2015.06.002.

43. Chin, A.F., D. Toptygin, W.A. Elam, T.P. Schrank, and V.J. Hilser. 2016. Phosphorylation Increases Persistence Length and End-to-End Distance of a Segment of Tau Protein. Biophys. J. 110:362–371, doi: 10.1016/j.bpj.2015.12.013.

44. Martin, E.W., A.S. Holehouse, C.R. Grace, A. Hughes, R.V. Pappu, and T. Mittag. 2016. Sequence Determinants of the Conformational Properties of an Intrinsically Disordered Protein Prior to and upon Multisite Phosphorylation. J. Am. Chem. Soc. 138:15323–15335, doi: 10.1021/jacs.6b10272.

45. Zeng, Y., D. Gao, J.J. Kim, T. Shiraishi, N. Terada, Y. Kakehi, C. Kong, R.H. Getzenberg, and P. Kulkarni. 2013. Prostate-associated gene 4 (PAGE4) protects cells against stress by elevating p21 and suppressing reactive oxygen species production. Am J Clin Exp Urol. 1:39–52.

46. Kulkarni, P., M.K. Jolly, D. Jia, S.M. Mooney, A. Bhargava, L.T. Kagohara, Y. Chen, P. Hao, Y. He, R.W. Veltri, A. Grishaev, K. Weninger, H. Levine, and J. Orban. 2017. Phosphorylation-induced conformational dynamics in an intrinsically disordered protein and potential role in phenotypic heterogeneity. Proc. Natl. Acad. Sci. U. S. A. 114:E2644– E2653, doi: 10.1073/pnas.1700082114.

47. He, Y., Y. Chen, S.M. Mooney, K. Rajagopalan, A. Bhargava, E. Sacho, K. Weninger, P.N. Bryan, P. Kulkarni, and J. Orban. 2015. Phosphorylation-induced Conformational Ensemble Switching in an Intrinsically Disordered Cancer/Testis Antigen. J. Biol. Chem. 290:25090– 25102, doi: 10.1074/jbc.M115.658583.

48. Lin, X., P. Kulkarni, F. Bocci, N.P. Schafer, S. Roy, M.Y. Tsai, Y. He, Y. Chen, K. Rajagopalan, S.M. Mooney, Y. Zeng, K. Weninger, A. Grishaev, J.N. Onuchic, H. Levine, P.G. Wolynes, R. Salgia, G. Rangarajan, V. Uversky, J. Orban, and M.K. Jolly. 2019. Structural and Dynamical Order of a Disordered Protein: Molecular Insights into Conformational Switching of PAGE4 at the Systems Level. Biomolecules. 9doi: 10.3390/biom9020077.

49. Vymětal, J., V. Jurásková, and J. Vondrášek. 2019. AMBER and CHARMM Force Fields Inconsistently Portray the Microscopic Details of Phosphorylation. J. Chem. Theory Comput. 15:665–679, doi: 10.1021/acs.jctc.8b00715.

50. Vitalis, A., and R.V. Pappu. 2009. ABSINTH: a new continuum solvation model for simulations of polypeptides in aqueous solutions. J. Comput. Chem. 30:673–699, doi: 10.1002/jcc.21005.

51. Vitalis, A., and R.V. Pappu. 2009. Chapter 3 Methods for Monte Carlo Simulations of Biomacromolecules. In: Annual Reports in Computational Chemistry. pp. 49–76, doi: 10.1016/s1574-1400(09)00503-9.

52. Peran, I., A.S. Holehouse, I.S. Carrico, R.V. Pappu, O. Bilsel, and D.P. Raleigh. 2019. Unfolded states under folding conditions accommodate sequence-specific conformational preferences with random coil-like dimensions. Proc. Natl. Acad. Sci. U. S. A. 116:12301– 12310, doi: 10.1073/pnas.1818206116.

53. Cook, E.C., D. Sahu, M. Bastidas, and S.A. Showalter. 2019. Solution Ensemble of the C-Terminal Domain from the Transcription Factor Pdx1 Resembles an Excluded Volume Polymer. J. Phys. Chem. B. 123:106–116, doi: 10.1021/acs.jpcb.8b10051.

54. Metskas, L.A., and E. Rhoades. 2015. Conformation and Dynamics of the Troponin I C-Terminal Domain: Combining Single-Molecule and Computational Approaches for a Disordered Protein Region. J. Am. Chem. Soc. 137:11962–11969, doi: 10.1021/jacs.5b04471.

55. Gibbs, E.B., and S.A. Showalter. 2016. Quantification of Compactness and Local Order in the Ensemble of the Intrinsically Disordered Protein FCP1. J. Phys. Chem. B. 120:8960– 8969, doi: 10.1021/acs.jpcb.6b06934.

56. Fuertes, G., N. Banterle, K.M. Ruff, A. Chowdhury, D. Mercadante, C. Koehler, M. Kachala, G. Estrada Girona, S. Milles, A. Mishra, P.R. Onck, F. Grater, S. Esteban-Martin, R.V. Pappu, D.I. Svergun, and E.A. Lemke. 2017. Decoupling of size and shape fluctuations in heteropolymeric sequences reconciles discrepancies in SAXS vs. FRET measurements. Proc. Natl. Acad. Sci. U. S. A. 114:E6342–E6351, doi: 10.1073/pnas.1704692114.

57. Das, R.K., Y. Huang, A.H. Phillips, R.W. Kriwacki, and R.V. Pappu. 2016. Cryptic sequence features within the disordered protein p27^Kip1^ regulate cell cycle signaling. Proc. Natl. Acad. Sci. U. S. A. 113:5616–5621, doi: 10.1073/pnas.1516277113.

58. Sherry, K.P., R.K. Das, R.V. Pappu, and D. Barrick. 2017. Control of transcriptional activity by design of charge patterning in the intrinsically disordered RAM region of the Notch receptor. Proc. Natl. Acad. Sci. U. S. A. 114:E9243–E9252, doi: 10.1073/pnas.1706083114.

59. Fossat, M.J., X. Zeng, and R.V. Pappu. 2021. Uncovering Differences in Hydration Free Energies and Structures for Model Compound Mimics of Charged Side Chains of Amino Acids. J. Phys. Chem. B. 125:4148–4161, doi: 10.1021/acs.jpcb.1c01073.

60. Moser, A., K. Range, and D.M. York. 2010. Accurate proton affinity and gas-phase basicity values for molecules important in biocatalysis. J Phys Chem B. 114:13911–13921, doi: 10.1021/jp107450n.

61. Platzer, G., M. Okon, and L.P. McIntosh. 2014. pH-dependent random coil (1)H, (13)C, and (15)N chemical shifts of the ionizable amino acids: a guide for protein pK a measurements. J. Biomol. NMR. 60:109–129, doi: 10.1007/s10858-014-9862-y.

62. Stull, D.R. 1947. Vapor Pressure of Pure Substances. Organic and Inorganic Compounds. Ind Eng Chem. 39:517–540, doi: 10.1021/ie50448a022.

63. Kumler, W.D., and J.J. Eiler. 1943. The Acid Strength of Mono and Diesters of Phosphoric Acid. The n-Alkyl Esters from Methyl to Butyl, the Esters of Biological Importance, and the Natural Guanidine Phosphoric Acids. J. Am. Chem. Soc. 65:2355–2361, doi: 10.1021/ja01252a028.

64. Dodda, L.S., I. Cabeza de Vaca, J. Tirado-Rives, and W.L. Jorgensen. 2017. LigParGen web server: an automatic OPLS-AA parameter generator for organic ligands. Nucleic Acids Res. 45:W331–W336, doi: 10.1093/nar/gkx312.

65. McGibbon, R.T., K.A. Beauchamp, M.P. Harrigan, C. Klein, J.M. Swails, C.X. Hernández, C.R. Schwantes, L.-P. Wang, T.J. Lane, and V.S. Pande. 2015. MDTraj: A Modern Open Library for the Analysis of Molecular Dynamics Trajectories. Biophys. J. 109:1528–1532, doi: 10.1016/j.bpj.2015.08.015.

66. Mao, A.H., and R.V. Pappu. 2012. Crystal lattice properties fully determine short-range interaction parameters for alkali and halide ions. J. Chem. Phys. 137:064104, doi: 10.1063/1.4742068.

67. Lalmansingh, J.M., A.T. Keeley, K.M. Ruff, R.V. Pappu, and A.S. Holehouse. 2023. SOURSOP: A Python Package for the Analysis of Simulations of Intrinsically Disordered Proteins. J. Chem. Theory Comput. 19:5609–5620, doi: 10.1021/acs.jctc.3c00190.

68. Humphrey, W., A. Dalke, and K. Schulten. 1996. VMD: visual molecular dynamics. J. Mol. Graph. 14:33–8, 27–8, doi: 10.1016/0263-7855(96)00018-5.

69. Pettersen, E.F., T.D. Goddard, C.C. Huang, E.C. Meng, G.S. Couch, T.I. Croll, J.H. Morris, and T.E. Ferrin. 2021. UCSF ChimeraX: Structure visualization for researchers, educators, and developers. Protein Sci. 30:70–82, doi: 10.1002/pro.3943.

70. Thirumalai, D., and B.-Y. Ha. 1997. Statistical Mechanics of Semiflexible Chains: A Meanfield Variational Approach. In: Grosberg A, editor. Theoretical and Mathematical Models in Polymer Research. Academic Press.

71. Dill, K.A., and S. Bromberg. 2011. Molecular Driving Forces: Statistical Thermodynamics in Biology, Chemistry, Physics, and Nanoscience. 2nd ed. Garland Science, Taylor & Francis Group, LLC.

72. Gibbs, E.B., F. Lu, B. Portz, M.J. Fisher, B.P. Medellin, T.N. Laremore, Y.J. Zhang, D.S. Gilmour, and S.A. Showalter. 2017. Phosphorylation induces sequence-specific conformational switches in the RNA polymerase II C-terminal domain. Nat. Commun. 8:15233, doi: 10.1038/ncomms15233.

73. Minor, D.L., Jr, and P.S. Kim. 1994. Measurement of the beta-sheet-forming propensities of amino acids. Nature. 367:660–663, doi: 10.1038/367660a0.

74. Pandey, A.K., H.K. Ganguly, S.K. Sinha, K.E. Daniels, G.P.A. Yap, S. Patel, and N.J. Zondlo. 2023. An Inherent Difference between Serine and Threonine Phosphorylation: Phosphothreonine Strongly Prefers a Highly Ordered, Compact, Cyclic Conformation. ACS Chem. Biol.doi: 10.1021/acschembio.3c00068.

75. Smart, J.L., and J.A. McCammon. 1999. Phosphorylation stabilizes the N-termini of alpha-helices. Biopolymers. 49:225–233, doi: 10.1002/(SICI)1097-0282(199903)49:3<225::AID-BIP4>3.0.CO;2-B.

76. Aurora, R., and G.D. Rose. 1998. Helix capping. Protein Sci. 7:21–38, doi: 10.1002/pro.5560070103.

77. Forood, B., E.J. Feliciano, and K.P. Nambiar. 1993. Stabilization of alpha-helical structures in short peptides via end capping. Proc. Natl. Acad. Sci. U. S. A. 90:838–842, doi: 10.1073/pnas.90.3.838.

78. Andrew, C.D., J. Warwicker, G.R. Jones, and A.J. Doig. 2002. Effect of phosphorylation on alpha-helix stability as a function of position. Biochemistry. 41:1897–1905, doi: 10.1021/bi0113216.

79. Kabsch, W., and C. Sander. 1983. Dictionary of protein secondary structure: pattern recognition of hydrogen-bonded and geometrical features. Biopolymers. 22:2577–2637, doi: 10.1002/bip.360221211.

80. Mooney, S.M., R. Qiu, J.J. Kim, E.J. Sacho, K. Rajagopalan, D. Johng, T. Shiraishi, P. Kulkarni, and K.R. Weninger. 2014. Cancer/testis antigen PAGE4, a regulator of c-Jun transactivation, is phosphorylated by homeodomain-interacting protein kinase 1, a component of the stress-response pathway. Biochemistry. 53:1670–1679, doi: 10.1021/bi500013w.

81. Brister, M.A., A.K. Pandey, A.A. Bielska, and N.J. Zondlo. 2014. OGlcNAcylation and phosphorylation have opposing structural effects in tau: phosphothreonine induces particular conformational order. J. Am. Chem. Soc. 136:3803–3816, doi: 10.1021/ja407156m.

82. Palumbo, P., M. Vanoni, V. Cusimano, S. Busti, F. Marano, C. Manes, and L. Alberghina. 2016. Whi5 phosphorylation embedded in the G1/S network dynamically controls critical cell size and cell fate. Nat. Commun. 7:11372, doi: 10.1038/ncomms11372.

83. Monahan, Z., V.H. Ryan, A.M. Janke, K.A. Burke, S.N. Rhoads, G.H. Zerze, R. O’Meally, G.L. Dignon, A.E. Conicella, W. Zheng, R.B. Best, R.N. Cole, J. Mittal, F. Shewmaker, and N.L. Fawzi. 2017. Phosphorylation of the FUS low-complexity domain disrupts phase separation, aggregation, and toxicity. EMBO J. 36:2951–2967, doi: 10.15252/embj.201696394.

84. Gruijs da Silva, L.A., F. Simonetti, S. Hutten, H. Riemenschneider, E.L. Sternburg, L.M. Pietrek, J. Gebel, V. Dötsch, D. Edbauer, G. Hummer, L.S. Stelzl, and D. Dormann. 2022. Disease-linked TDP-43 hyperphosphorylation suppresses TDP-43 condensation and aggregation. EMBO J. 41:e108443, doi: 10.15252/embj.2021108443.

85. Yamazaki, H., M. Takagi, H. Kosako, T. Hirano, and S.H. Yoshimura. 2022. Cell cycle-specific phase separation regulated by protein charge blockiness. Nat. Cell Biol. 24:625– 632, doi: 10.1038/s41556-022-00903-1.

86. Paleologou, K.E., A.W. Schmid, C.C. Rospigliosi, H.-Y. Kim, G.R. Lamberto, R.A. Fredenburg, P.T. Lansbury, C.O. Fernandez, D. Eliezer, M. Zweckstetter, and H.A. Lashuel. 2008. Phosphorylation at Ser-129 but Not the Phosphomimics S129E/D Inhibits the Fibrillation of α-Synuclein*. J. Biol. Chem. 283:16895–16905, doi: 10.1074/jbc.M800747200.

87. Somale, D., G. Di Nardo, L. di Blasio, A. Puliafito, M. Vara-Messler, G. Chiaverina, M. Palmiero, V. Monica, G. Gilardi, L. Primo, and P.A. Gagliardi. 2020. Activation of RSK by phosphomimetic substitution in the activation loop is prevented by structural constraints. Sci. Rep. 10:591, doi: 10.1038/s41598-019-56937-3.

88. Kozeleková, A., A. Náplavová, T. Brom, N. Gašparik, J. Šimek, J. Houser, and J. Hritz. 2022. Phosphorylated and Phosphomimicking Variants May Differ-A Case Study of 14-3-3 Protein. Front Chem. 10:835733, doi: 10.3389/fchem.2022.835733.

89. Dong, X., Z. Gong, Y.B. Lu, K. Liu, L.Y. Qin, M.L. Ran, C.L. Zhang, Z. Liu, W.P. Zhang, and C. Tang. 2017. Ubiquitin S65 phosphorylation engenders a pH-sensitive conformational switch. Proc. Natl. Acad. Sci. U. S. A. 114:6770–6775, doi: 10.1073/pnas.1705718114.

90. Lei, R., J.P. Lee, M.B. Francis, and S. Kumar. 2018. Structural Regulation of a Neurofilament-Inspired Intrinsically Disordered Protein Brush by Multisite Phosphorylation. Biochemistry. 57:4019–4028, doi: 10.1021/acs.biochem.8b00007.

91. Malka-Gibor, E., M. Kornreich, A. Laser-Azogui, O. Doron, I. Zingerman-Koladko, J. Harapin, O. Medalia, and R. Beck. 2017. Phosphorylation-Induced Mechanical Regulation of Intrinsically Disordered Neurofilament Proteins. Biophys. J. 112:892–900, doi: 10.1016/j.bpj.2016.12.050.

92. Marchand, J.-R., T. Knehans, A. Caflisch, and A. Vitalis. 2020. An ABSINTH-Based Protocol for Predicting Binding Affinities between Proteins and Small Molecules. J. Chem. Inf. Model. 60:5188–5202, doi: 10.1021/acs.jcim.0c00558.

93. Fossat, M.J., and R.V. Pappu. 2019. q-Canonical Monte Carlo Sampling for Modeling the Linkage between Charge Regulation and Conformational Equilibria of Peptides. J. Phys. Chem. B. 123:6952–6967, doi: 10.1021/acs.jpcb.9b05206.

94. Fossat, M.J., A.E. Posey, and R.V. Pappu. 2021. Quantifying charge state heterogeneity for proteins with multiple ionizable residues. Biophys. J. 120:5438–5453, doi: 10.1016/j.bpj.2021.11.2886.

95. Fossat, M.J., A.E. Posey, and R.V. Pappu. 2023. Uncovering the Contributions of Charge Regulation to the Stability of Single Alpha Helices. Chemphyschem. 24:e202200746, doi: 10.1002/cphc.202200746.

96. Crabtree, M.D., J. Holland, A.S. Pillai, P.S. Kompella, L. Babl, N.N. Turner, J.T. Eaton, G.K.A. Hochberg, D.G.A.L. Aarts, C. Redfield, A.J. Baldwin, and T.J. Nott. 2023. Ion binding with charge inversion combined with screening modulates DEAD box helicase phase transitions. Cell Rep. 42:113375, doi: 10.1016/j.celrep.2023.113375.

