## Supplementary Information for "Phosphorylation of disordered proteins tunes local and global intramolecular interactions"

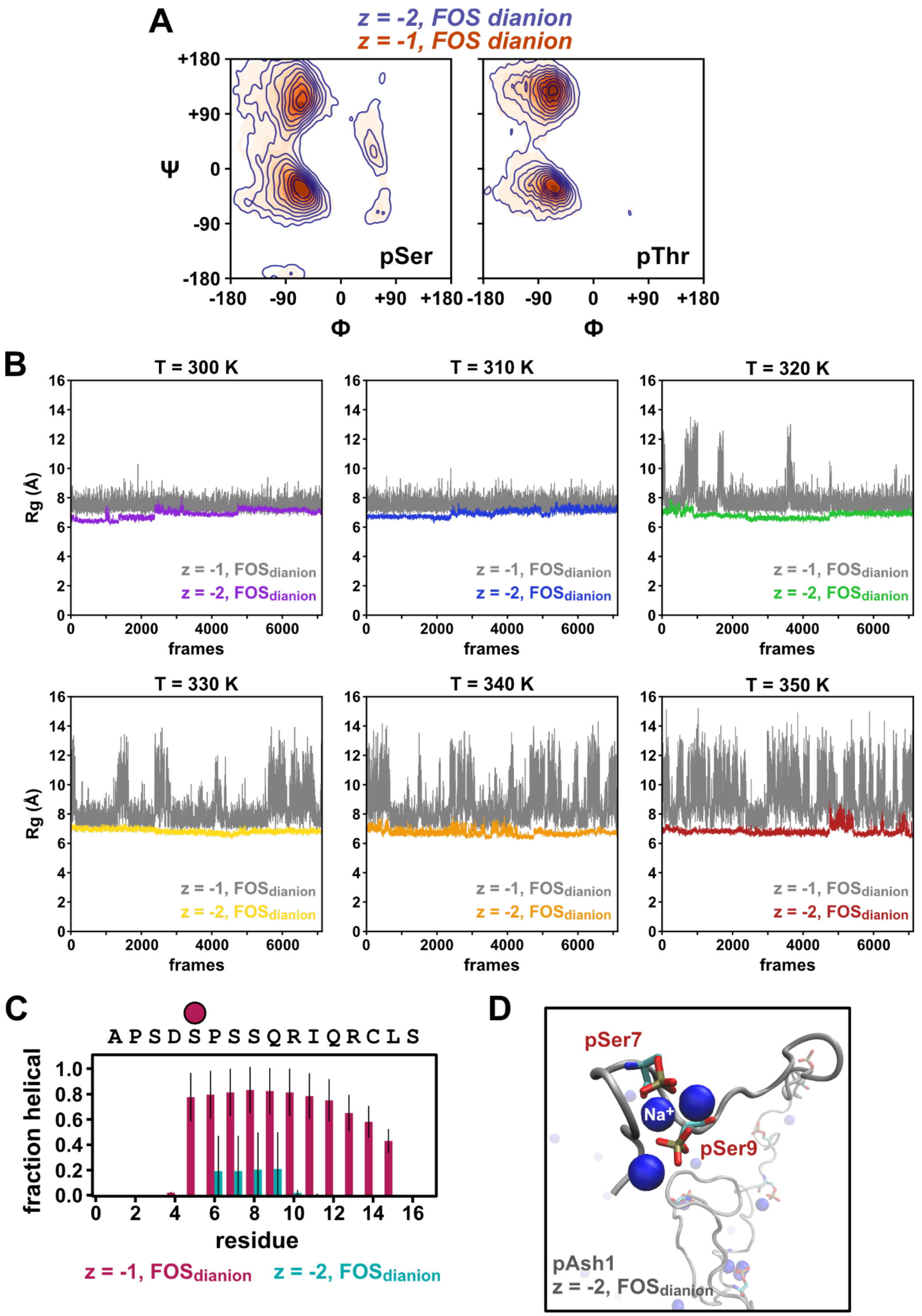


**Figure S1: Comparison of sampling quality between z = -1 and z = -2 parameters. (A)** Ramachandran plots for G-pS-G and G-pT-G simulations showing good agreement of the torsion angle space between z = -1 and z = -2 (both using free energy of solvation for a dianion ($FOS_{dianion}$)). **(B)** Radius of gyration per frame in pTH4 simulations at various temperatures for z

= -1 with FOS$_{dianion}$ (gray traces) and z = -2 with FOS$_{dianion}$ (colored traces). A comparison of the traces suggests that the pTH4 peptide in z = -2 simulations is trapped in a collapsed conformation, even at higher temperatures. **(C)** Plot of helical content per residue for pTH4 in z = -1 (magenta) or z = -2 (cyan) simulations. Error bars represent the standard deviation from the mean values determined from each of the replicates. The magenta circle denotes the site of phosphorylation. **(D)** Snapshot of pAsh1 simulation using z = -2, FOS$_{dianion}$ parameters. pSer7 and pSer9 appear chelate sodium ions using the phosphoryl group and carbonyl oxygen of the backbone. This behavior leads to poor sampling because the sodium ions are relatively stably bound.

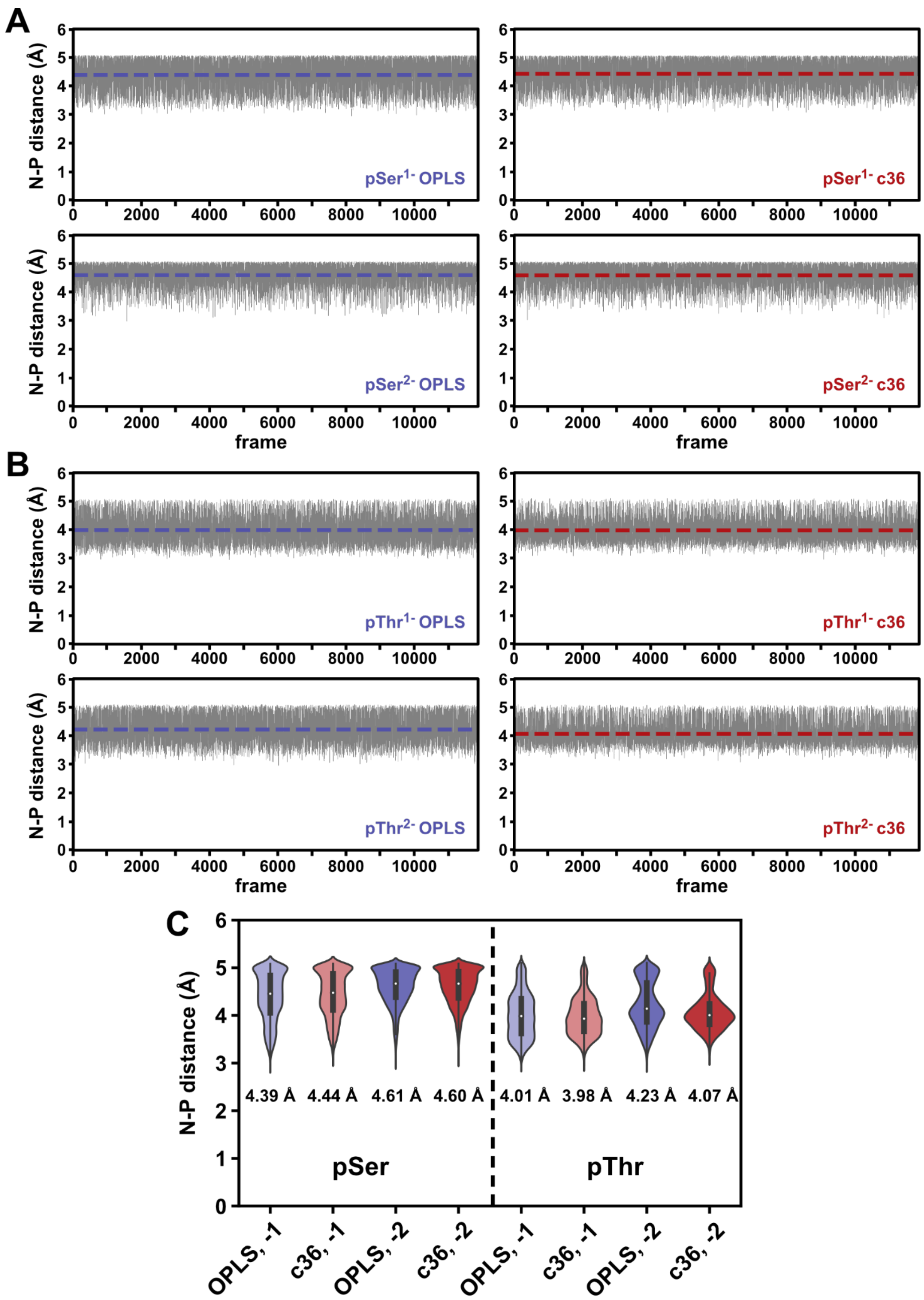


**Figure S2: Modest differences in local conformational space between pSer and pThr. (A)** and **(B)** Plots of the N-P distances (the distance from the phosphoryl phosphorus to the amide nitrogen of the phosphorylated residue) from G-pS-G or G-pT-G simulations over the course of the simulation. The average distance is plotted as a horizontal dashed line. For a given phospho-residue and charge state, the OPLS (blue) and CHARMM36 (c36) (red) forcefield parameters yield very similar results. The "-2" results presented in this panel were generated from simulations using z = -1, $FOS_{dianion}$ parameters. **(C)** Summary of phosphoryl-amide distances. Average distances are shown below each violin.

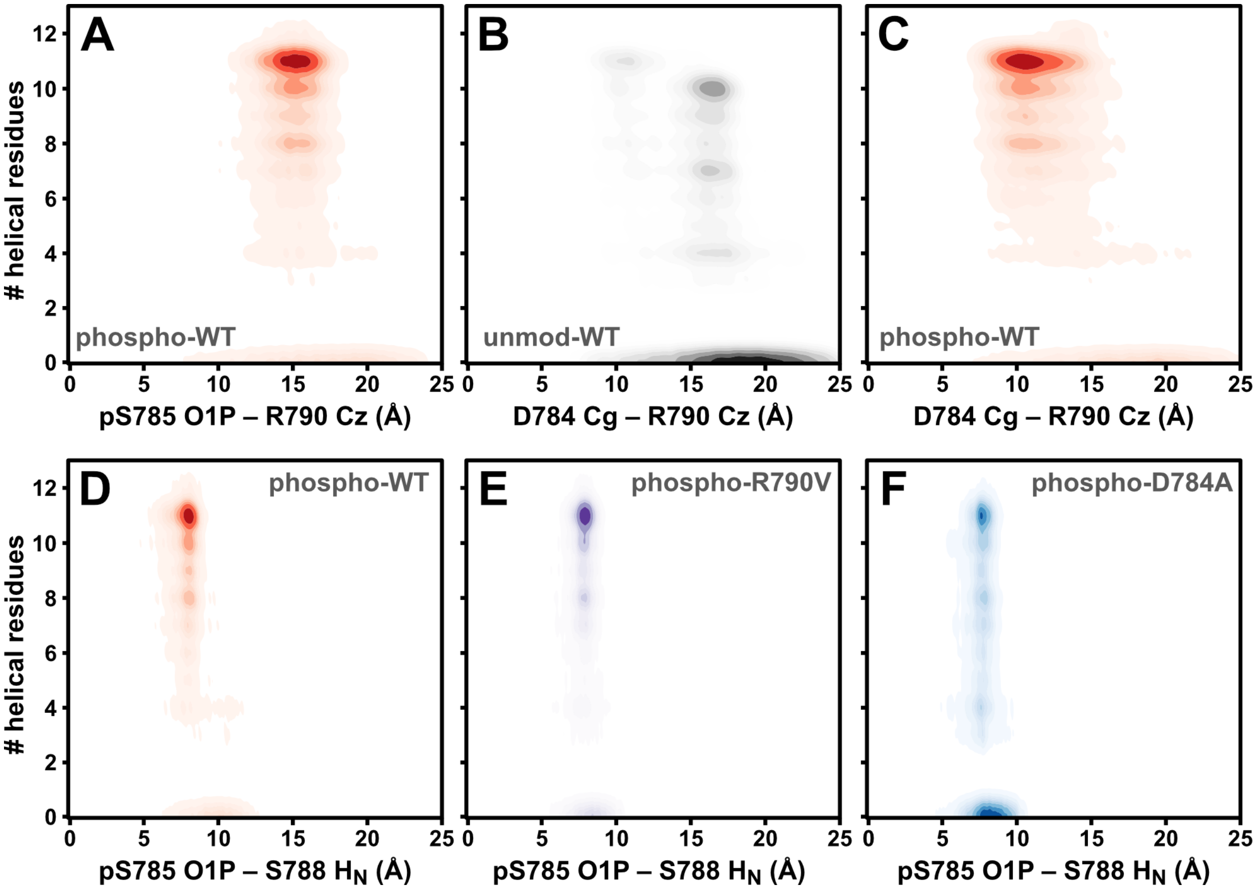


**Figure S3: Insights into the interactions that stabilize the TH4 peptide helix in the phosphorylated state. (A)** 2D distributions showing the number of helical residues plotted against the distance between the phosphoryl group (residue i) to the i+5 Arg side chain. **(B)** Number of helical residues versus the distance between the i-1 Asp and i+5 Arg in the unmodified and **(C)** pTH4 simulations. Comparison of panels **(B)** and **(C)** highlights the closer proximity of the i-1 and i+5 side chains upon phosphorylation. **(D)** through **(F)** Number of helical residues versus the distance between the phosphoryl group and the backbone of the i+3 residue for **(D)** WT, **(E)** R790V, and **(F)** D784A pTH4 peptides.

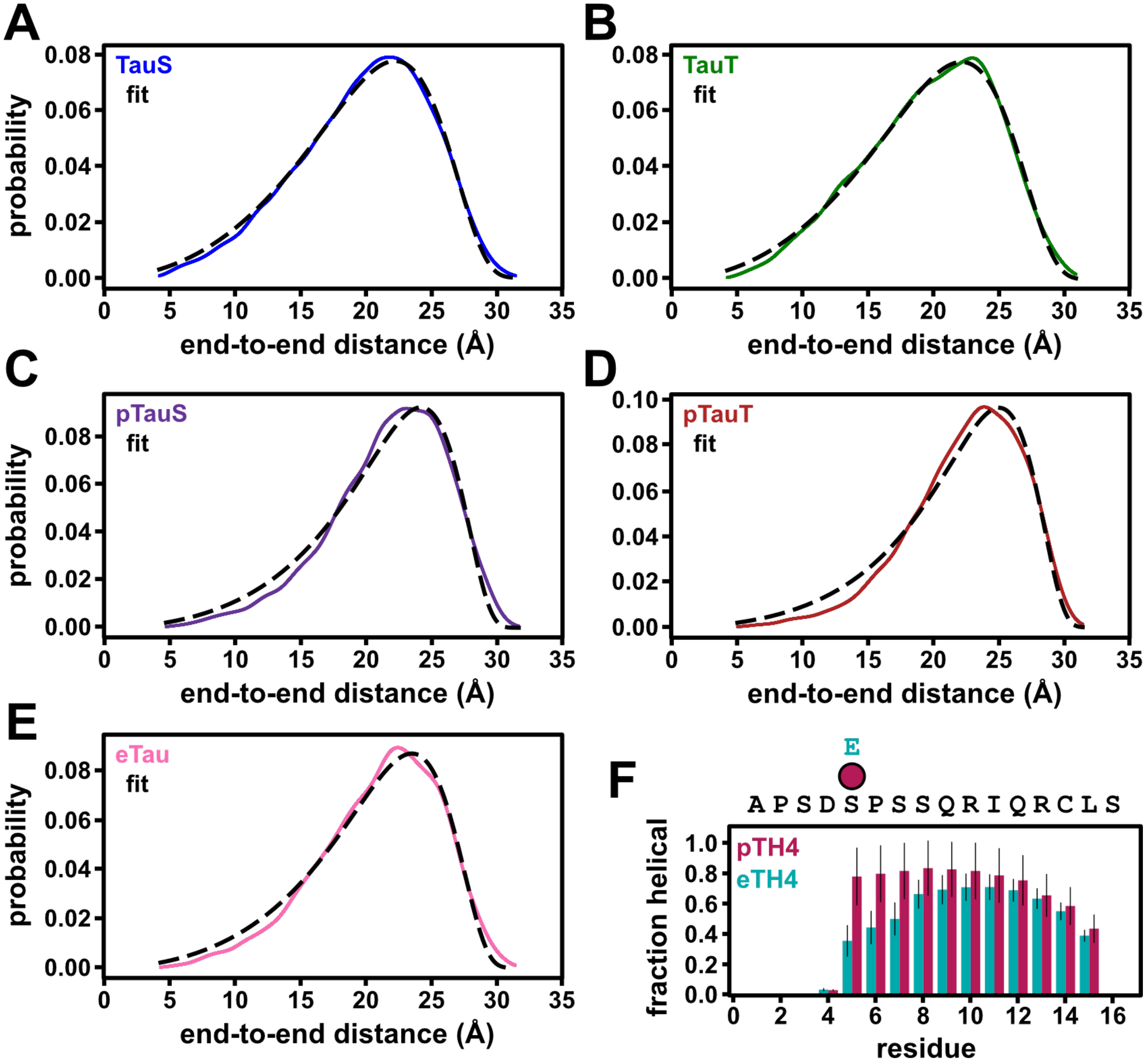


**Figure S4: Ensemble features of short phosphopeptides**. **(A)** through **(E)** End-to-end distance distributions (colors) and nonlinear fits for persistence length (dashed lines) for **(A)** TauS, **(B)** TauT, **(C)** pTauS, **(D)** pTauT, and **(E)** eTau. **(F)** Comparison of helical content per residue for pTH4 and eTH4. Error bars represent the standard deviation from the mean values determined from each of the replicates.

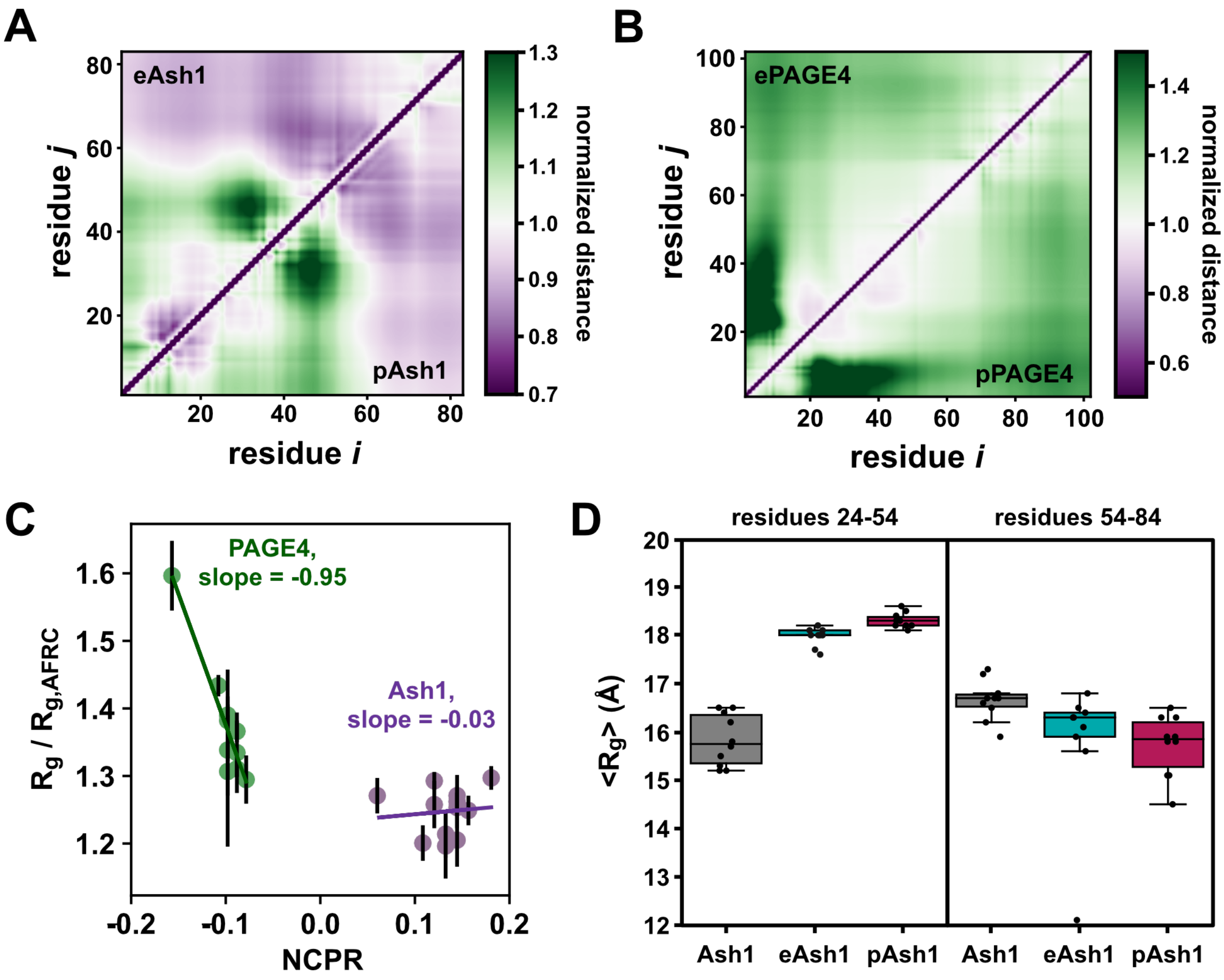


**Figure S5: Phosphorylation induced ensemble effects on long IDRs with multiple phosphorylation events. (A)** Distance maps of phosphomimetic Ash1 (eAsh1) (top left) or pAsh1 (bottom left) normalized against the unmodified Ash1. **(B)** Distance maps of phosphomimetic PAGE4 (ePAGE4) (top left) or pPAGE4 (bottom left) normalized against the unmodified PAGE4. **(C)** Ensemble dimensions (normalized against the dimensions of a random coil of the same length) for PAGE4 and Ash1 as a function of net charge per residue (NCPR) for each phosphoform. **(D)** Ensemble dimensions for sub-regions of Ash1, eAsh1, and pAsh1.

**Table S1: ABSINTH-OPLS/AA parameters for phosphorylated Ser and Thr implemented in this study.**

| atom | # occurrences | charge | charge x occurrences | net charge | FOS (free energy of solvation) (kJ/mol) |
|---|---|---|---|---|---|
| **phosphoserine** (pS / pSer / Sep) | | | | | |
| Cb | 1 | -0.071 | -0.071 | | |
| Og | 1 | -0.798 | -0.798 | | |
| P | 1 | 2.761 | 2.761 | | |
| O-P | 1 | -0.986 | -0.986 | -1* | -96.8 |
| O=P | 2 | -1.227 | -2.454 | | |
| HO-P | 1 | 0.396 | 0.396 | | |
| Hb | 2 | 0.076 | 0.152 | | |
| Cb | 1 | 0.092 | 0.092 | | |
| Og | 1 | -0.822 | -0.822 | | |
| P | 1 | 2.475 | 2.475 | -2 | -289.6* |
| O-P | 1 | -1.299 | -1.299 | | |
| O=P | 2 | -1.299 | -2.598 | | |
| Hb | 2 | 0.076 | 0.152 | | |
| **phosphothreonine** (pT / pThr / Tpo) | | | | | |
| Cb | 1 | 0.138 | 0.138 | | |
| Og | 1 | -0.816 | -0.816 | -1[§] | -96.8 |
| Cg | 1 | -0.180 | -0.180 | | |

| | | | | | |
|---|---|---|---|---|---|
| Hg | 3 | 0.06 | 0.180 | | |
| P | 1 | 2.621 | 2.621 | | |
| O-P | 1 | -0.990 | -0.990 | | |
| O=P | 2 | -1.221 | -2.442 | | |
| HO-P | 1 | 0.396 | 0.396 | | |
| Hb | 1 | 0.094 | 0.094 | | |
| Cb | 1 | 0.138 | 0.138 | -2 | -289.1§ |
| Og | 1 | -0.816 | -0.816 | | |
| Cg | 1 | -0.18 | -0.18 | | |
| Hg | 3 | 0.06 | 0.18 | | |
| P | 1 | 2.478 | 2.478 | | |
| O-P | 1 | -1.298 | -1.298 | | |
| O=P | 2 | -1.298 | -2.596 | | |
| Hb | 1 | 0.094 | 0.094 | | |

Underlines in atom identifiers denote which atom harbors the parameters in that row. The paired special symbols (* and §) for each pSer and pThr are provided to show the combination of charge and solvation parameters used in this work that yielded well-balanced charge interactions and well-sampled trajectories.

**Table S2: Details of CAMPARI simulations for all variants of the given system**

| system | temperature (K) | [NaCl] (M) | # equilibration frames / rep. | # production frames / rep. | replicates | total # frames |
|---|---|---|---|---|---|---|
| G-pS/T-G | 300 | 0.05 | 2,500,000 | 50,000,000 | 5 | 11,785 |
| TH4 | 330 | 0.15 | 2,500,000 | 50,000,000 | 10* | 23,570 |
| Tau | 300 | 0 | 2,500,000 | 50,000,000 | 5 | 11,785 |
| Ash1 | 320 | 0.05 | 2,500,000 | 50,000,000 | 10 | 23,570 |
| PAGE4 | 320 | 0.1 | 2,500,000 | 50,000,000 | 10 | 23,570 |

*TH4 simulations were performed using both coil and helical starting conformations (five independent replicates of each)

**Table S3: Comparison of Tau peptide persistence lengths from published experiments and our simulations.**

| | published[a] | | this work[b] | | |
|---|---|---|---|---|---|
| peptide | $\langle R_{ee} \rangle$ (Å) | $L_P$ (Å) | $\langle R_{ee} \rangle$ (Å) | $L_P$ (Å) | $L_P$ (residues) |
| TauT | 29.445 (0.065) | 12.487 (0.075) | 19.8 (0.1) | 7.38 (0.01) | 2.63 (0.01) |
| TauS | 29.173 (0.047) | 12.18 (0.053) | 19.8 (0.1) | 7.48 (0.01) | 2.64 (0.01) |
| pTauT | 31.689 (0.038) | 15.349 (0.054) | 22.5 (0.1) | 10.75 (0.01) | 4.55 (0.04) |
| pTauS | 31.000 (0.035) | 14.401 (0.046) | 21.7 (0.2) | 9.79 (0.01) | 3.86 (0.01) |
| eTau | ND | ND | 21.0 (0.2) | 8.98 (0.01) | 3.42 (0.02) |

[a]published Tau measurements were taken from (1); the error reported in parentheses is the 95% confidence interval

[b]Tau distances and persistence lengths from our simulations; the error reported in parentheses is the standard error of fit determined from the covariance matrix

ND = no data reported

**Table S4: Ash1 phospho-states and ensemble dimensions from simulations.** All Ash1 phospho-state simulations were conducted as described in **Table S2.**

| phospho-protein | # phosphates | sequence (phosphorylated sites) | NCPR* | $\langle R_G \rangle$ (Å)§ |
|---|---|---|---|---|
| Ash1 | 0 | GASASSSPSPSTPTKSGKMRSRSSSP<br>VRPKAYTPSPRSPNYHRFALDSPPQS<br>PRRSSNSSITKKGSRRSSGSSPTRHTT<br>RVCV | 0.1807 | 29.6 (0.4) |
| pAsh1 | 10 | GASASS**S**P**S**PS**T**PTKSGKMRSRSS**S**P<br>VRPKAY**T**P**S**PR**S**PNYHRFALD**S**PPQ**S**<br>PRRSSNSSITKKGSRRSSGS**S**PTRHTT<br>RVCV | 0.0602 | 29.0 (0.6) |
| eAsh1 | 0 (10x Glu) | GASASS**E**P**E**PS**E**PTKSGKMRSRSS**E**P<br>VRPKAY**E**P**E**PR**E**PNYHRFALD**E**PPQ**E**<br>PRRSSNSSITKKGSRRSSGS**E**PTRHTT<br>RVCV | 0.1807 | 28.4 (1.1) |
| p[3x]-Ash1 | 3 | GASASS**S**P**S**PS**T**PTKSGKMRSRSSSP<br>VRPKAYTPSPRSPNYHRFALDSPPQS<br>PRRSSNSSITKKGSRRSSGSSPTRHTT<br>RVCV | 0.1446 | 27.4 (0.9) |
| p[4x]-Ash1 | 4 | GASASS**S**P**S**PS**T**PTKSGKMRSRSS**S**P<br>VRPKAYTPSPRSPNYHRFALDSPPQS<br>PRRSSNSSITKKGSRRSSGSSPTRHTT<br>RVCV | 0.1325 | 27.3 (1.1) |
| p[5xA]-Ash1 | 5 | GASASS**S**PSPS**T**PTKSGKMRSRSSSP<br>VRPKAY**T**PSPR**S**PNYHRFALDSPPQ**S**<br>PRRSSNSSITKKGSRRSSGSSPTRHTT<br>RVCV | 0.1205 | 28.6 (0.8) |
| p[5xB]-Ash1 | 5 | GASASSSP**S**PSTPTKSGKMRSRSS**S**P<br>VRPKAYTP**S**PRSPNYHRFALD**S**PPQS<br>PRRSSNSSITKKGSRRSSGS**S**PTRHTT<br>RVCV | 0.1205 | 29.6 (0.4) |
| seq13† | 3 | GASASSSPSPSTPTKSGKMRSRSSSP<br>VRPKAY**T**P**S**PR**S**PNYHRFALDSPPQS<br>PRRSSNSSITKKGSRRSSGSSPTRHTT<br>RVCV | 0.1446 | 27.5 (0.9) |
| seq14† | 5 | GASASS**S**P**S**PS**T**PTKSGKMRSRSSSP<br>VRPKAY**T**P**S**PR**S**PNYHRFALDSPPQS<br>PRRSSNSSITKKGSRRSSGSSPTRHTT<br>RVCV | 0.1084 | 27.3 (1.1) |
| seq15† | 2 | GASASSSPSPSTPTKSGKMRSRSSSP<br>VRPKAYTPSPRSPNYHRFALD**S**PPQ**S**<br>PRRSSNSSITKKGSRRSSGSSPTRHTT<br>RVCV | 0.1566 | 28.5 (0.5) |

| | | | | |
|---|---|---|---|---|
| seq18† | 4 | GASASS**S**P**S**PS**T**PTKSGKMRSRSSSP<br>VRPKAYTPSPRSPNYHRFALDSPPQS<br>PRRSSNSSITKKGSRRSSGS**S**PTRHTT<br>RVCV | 0.1325 | 27.7 (0.8) |
| seq19† | 3 | GASASS**S**PSPS**T**PTKSGKMRSRSSSP<br>VRPKAYTP**S**PRSPNYHRFALDSPPQS<br>PRRSSNSSITKKGSRRSSGSSPTRHTT<br>RVCV | 0.1446 | 28.6 (0.7) |
| seq20† | 3 | GASASSSP**S**PSTPTKSGKMRSRSSSP<br>VRPKAY**T**PSPR**S**PNYHRFALDSPPQS<br>PRRSSNSSITKKGSRRSSGSSPTRHTT<br>RVCV | 0.1446 | 29.0 (0.7) |

§Ensemble-averaged radius of gyration (<$R_G$>); error in parentheses is the standard deviation from the mean of ten independent replicate simulations

*NCPR = net charge per residue for the given phospho-state calculated using CIDER (2); all values assume a phosphoryl group with z = -1

†phospho-Ash1 sequences that were not included in **Figure 5**, but were used to assess possible trends in $R_G$ as a function of NCPR in **Figure S5C**

**Table S5: PAGE4 phospho-states and ensemble dimensions from simulations.** All PAGE4 phospho-state simulations were conducted as described in **Table S2.**

| phospho-protein | # phosphates | sequence (phosphorylated sites) | NCPR* | $\langle R_G \rangle$ (Å)§ |
|---|---|---|---|---|
| PAGE4 | 0 | MSARVRSRSRGRGDGQEAPDVVAFV<br>APGESQQEEPPTDNQDIEPGQEREGT<br>PPIEERKVEGDCQEMDLEKTRSERGD<br>GSDVKEKTPPNPKHAKTKEAGDGQP | -0.0784 | 32.6 (0.9) |
| pPAGE4 | 8 | MSARVR**S**R**S**RGRGDGQEAPDVVAFV<br>APGESQQEEPPTDNQDIEPGQEREG**T**<br>PPIEERKVEGDCQEMDLEK**T**R**S**ERGD<br>G**S**DVKEK**T**PPNPKHAK**T**KEAGDGQP | -0.1569 | 40.2 (1.3) |
| ePAGE4 | 0 (8x Glu) | MSARVR**E**R**E**RGRGDGQEAPDVVAFV<br>APGESQQEEPPTDNQDIEPGQEREG**E**<br>PPIEERKVEGDCQEMDLEK**E**R**E**ERGD<br>G**E**DVKEK**E**PPNPKHAK**E**KEAGDGQP | -0.1569 | 40.0 (0.2) |
| p[2xN]-PAGE4 | 2 | MSARVR**S**R**S**RGRGDGQEAPDVVAFV<br>APGESQQEEPPTDNQDIEPGQEREGT<br>PPIEERKVEGDCQEMDLEKTRSERGD<br>GSDVKEKTPPNPKHAKTKEAGDGQP | -0.0980 | 35.0 (1.7) |
| p[2xC]-PAGE4 | 2 | MSARVRSRSRGRGDGQEAPDVVAFV<br>APGESQQEEPPTDNQDIEPGQEREGT<br>PPIEERKVEGDCQEMDLEK**T**R**S**ERGD<br>GSDVKEKTPPNPKHAKTKEAGDGQP | -0.0980 | 32.9 (2.8) |
| p[51]-PAGE4 | 1 | MSARVRSRSRGRGDGQEAPDVVAFV<br>APGESQQEEPPTDNQDIEPGQEREG**T**<br>PPIEERKVEGDCQEMDLEKTRSERGD<br>GSDVKEKTPPNPKHAKTKEAGDGQP | -0.0882 | 33.0 (0.3) |
| p[3x]-PAGE4 | 3 | MSARVR**S**R**S**RGRGDGQEAPDVVAFV<br>APGESQQEEPPTDNQDIEPGQEREG**T**<br>PPIEERKVEGDCQEMDLEKTRSERGD<br>GSDVKEKTPPNPKHAKTKEAGDGQP | -0.1078 | 36.1 (0.4) |
| seq4† | 1 | MSARVR**S**RSRGRGDGQEAPDVVAFV<br>APGESQQEEPPTDNQDIEPGQEREGT<br>PPIEERKVEGDCQEMDLEKTRSERGD<br>GSDVKEKTPPNPKHAKTKEAGDGQP | -0.0882 | 34.4 (0.4) |
| seq5† | 1 | MSARVRSR**S**RGRGDGQEAPDVVAFV<br>APGESQQEEPPTDNQDIEPGQEREGT<br>PPIEERKVEGDCQEMDLEKTRSERGD<br>GSDVKEKTPPNPKHAKTKEAGDGQP | -0.0882 | 33.6 (1.5) |
| seq8† | 2 | MSARVR**S**RSRGRGDGQEAPDVVAFV<br>APGESQQEEPPTDNQDIEPGQEREG**T**<br>PPIEERKVEGDCQEMDLEKTRSERGD<br>GSDVKEKTPPNPKHAKTKEAGDGQP | -0.0980 | 34.8 (0.2) |

| | | | | |
|---|---|---|---|---|
| seq9† | 2 | MSARVRSR**S**RGRGDGQEAPDVVAFV APGESQQEEPPTDNQDIEPGQEREG**T** PPIEERKVEGDCQEMDLEKTRSERGD GSDVKEKTPPNPKHAKTKEAGDGQP | -0.0980 | 33.7 (1.7) |

§Ensemble-averaged radius of gyration ($\langle R_G \rangle$); error in parentheses is the standard deviation from the mean of ten independent replicate simulations

*NCPR = net charge per residue for the given phospho-state calculated using CIDER (2)

†phospho-PAGE4 sequences that were not included in **Figure 5**, but were used to assess possible trends in $R_G$ as a function of NCPR in **Figure S5C**

### SUPPLEMENTAL REFERENCES

1. Chin AF, Toptygin D, Elam WA, Schrank TP, Hilser VJ. Phosphorylation Increases Persistence Length and End-to-End Distance of a Segment of Tau Protein. Biophys J. 2016;110(2):362–371. PMID: 26789759

2. Holehouse AS, Das RK, Ahad JN, Richardson MOG, Pappu RV. CIDER: Resources to Analyze Sequence-Ensemble Relationships of Intrinsically Disordered Proteins. Biophys J. 2017 Jan 10;112(1):16–21. PMID: 28076807
